# Selective inhibitory control of pyramidal neuron ensembles and cortical subnetworks by chandelier cells

**DOI:** 10.1101/140822

**Authors:** Jiangteng Lu, Jason Tucciarone, Nancy Padilla-Coreano, Miao He, Joshua A. Gordon, Z. Josh Huang

## Abstract

The neocortex comprises multiple information processing streams mediated by subsets of glutamatergic pyramidal cells (PCs) that receive diverse inputs and project to distinct targets. How GABAergic interneurons regulate the segregation and communication among intermingled PC subsets that contribute to separate brain networks remains unclear. Here we demonstrate that a subset of GABAergic chandelier cells (ChCs) in the prelimbic cortex (PL), which innervate PCs at spike initiation site, selectively control PCs projecting to the basolateral amygdala (_BLA_PC) compared to those projecting to contralateral cortex (_cc_PC). These ChCs in turn receive preferential input from local and contralateral _CC_PCs as opposed to _BLA_PCs and BLA neurons (the PL-BLA network). Accordingly, optogenetic activation of ChCs rapidly suppresses _BLA_PCs and BLA activity in freely behaving mice. Thus, the exquisite connectivity of ChCs not only mediates directional inhibition between local PC ensembles but may also shape communication hierarchies between global networks.

---

In many areas of the cerebral cortex, diverse and often intermingled subsets of pyramidal cells (PCs) preferentially receive inputs from and project outputs to distinct brain areas, and thus are embedded in separate local circuits as well as global networks^1^. It is not well understood how specific physiological PC ensembles emerge from the underlying anatomic scaffold and contribute to different subnetworks and information processing streams. Diverse types of GABAergic interneurons appear to specialize in their inhibitory control of various aspects of cortical circuit operations such as balancing excitation, modulating gain, tuning dynamics, and generating oscillations^2–4^. However, the inhibitory mechanisms that regulate the dynamic segregation of functional PC ensembles and route information flow between brain networks remain elusive.

Chandelier cells (ChCs, i.e. axo-axonic cells) are among the most distinctive interneuron types. ChCs selectively innervate PCs at their axon initial segment (AIS), the site of action potential initiation^5^. A single ChC innervates hundreds of PCs^6,7^, and multiple ChCs can converge onto the same PC^8,9^. The exquisite specificity of ChC innervation at AIS has long been speculated to exert the ultimate inhibitory control over PC spiking and population output^10,11^. However, it remains unclear how ChCs are recruited and whether a ChC non-discriminately innervates PCs within its dense axonal arbor or selects a specific PC subset^9^. In fact, it is even controversial whether ChCs inhibit or excite PCs^12–14^. Thus the problem of how ChCs control PCs represents a prominent gap as well as a unique opportunity for understanding the cellular basis of cortical organization, which entails elucidating the connectivity pattern of ChCs to PC subsets within local circuits in the context of global brain networks.

The rodent prelimbic area (PL) integrates inputs from the amygdala and other brain structures (e.g. other cortical areas, ventral hippocampus, medial-dorsal thalamus) to gate fear expression via projections back to the amygdala circuitry^15–19^. The superficial layers of PL contain two subsets of PCs: one projects to the basal lateral amygdala (BLAPC) and another that projects to contralateral cortex (_CC_PC)^15,20^. They form two separate subnetworks: the PL-BLA network, comprised of reciprocally connected _BLA_PCs and BLA neurons, and the bilateral PL network, comprised of _CC_PCs from the two hemispheres^20^. Here, by combining genetic labeling of ChCs and projection-based labeling of PC subsets, we demonstrate that a subset of layer 2 (L2) ChCs preferentially receives inputs from _CC_PCs yet selectively innervates _BLA_PCs. This highly directional ChC microcircuit module is distinct from the parvalbumin fast-spiking basket cell (PVBCs) module, characterized by non-selective and extensive reciprocal connectivity with _BLA_PCs and _CC_PCs. Trans-synaptic rabies tracing combined with optogenetic tagging of long-range inputs further revealed that L2 ChCs are preferentially recruited by contralateral _CC_PCs, but not by BLA input. Importantly, optogenetic activation of ChCs resulted in rapid inhibition of PCs firing in freely moving mice. Together, these results reveal that the exquisite connectivity of ChCs not only mediates directional inhibitory control between local PC ensembles but may also shape communication hierarchy and route information flow between distinct PC-associated global networks.

## RESULTS

### A subset of L2 ChCs selectively innervates _BLA_PCs over _CC_PCs in PL

We combined genetic^21^ and anatomic methods to reliably label ChCs, _BLA_PCs and _CC_PCs for physiological studies. Tamoxifen (TM) induction in pregnant *Nkx2.1-CreER;Rosa26-loxpSTOPloxp-TdTomato (Ai14)* mice at embryonic day 17.5 (E17.5) resulted in specific labeling of a subset of L2 ChCs throughout the frontal cortex, characterized by their somata position at the L1-L2 border, prominent dendritic arborization in L1, and dense axonal plexus in L2/3 (Fig. 1a, b and Supplementary Fig. 1). It should be noted that L2 ChCs are also generated at earlier embryonic times^21^; for simplicity the E17.5-born subset of L2 ChCs are herein referred to as “L2 ChCs”. Single cell reconstruction revealed that individual L2 ChCs elaborated on average 211±28 “cartridges”, vertical strings of boutons targeting the AIS of PCs (Fig. 1b and Supplementary Fig. 2). We distinguished subpopulations of L2/3 PCs in PL according to their projection targets by injecting different colors of retrograde tracer cholera toxin subunit B (CTB) into the BLA (to label _BLA_PCs), contralateral cortex (to label _CC_PCs) and dorsomedial striatum (to label _ST_PCs) of the same mouse (Fig.1c). Each PC population resided at characteristic laminar depths with some overlap; L2 ChCs occupied a similar laminar depth as _BLA_PCs (Fig. 1c, d and Supplementary Fig. 3a). Notably, there was little convergence in projection targets between _BLA_PCs and _CC_PCs (Fig. 1e).

**Figure 1.**
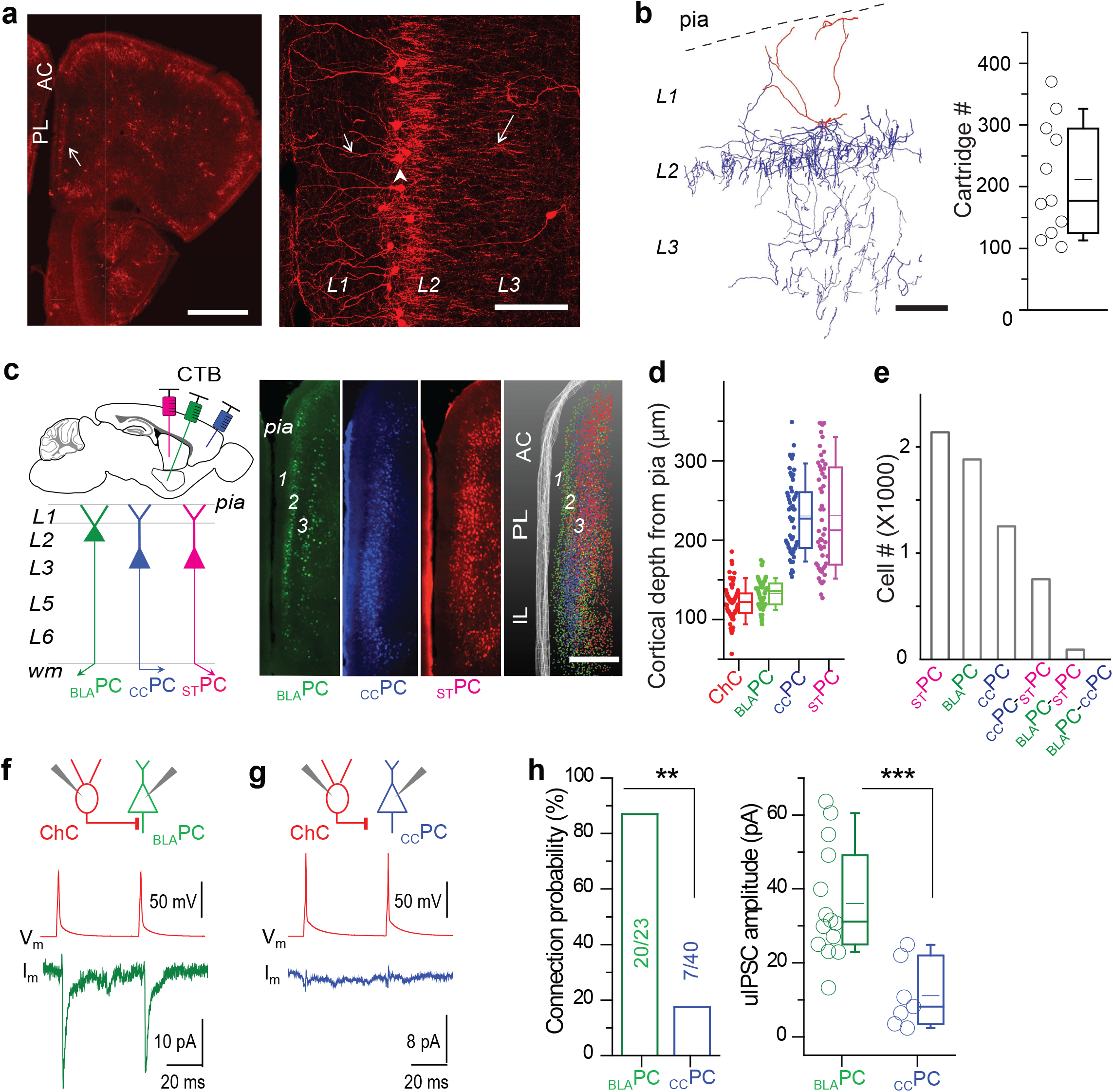
L2 ChCs preferentially innervate _BLA_PCs over _CC_PCs in prelimbic cortex. (**a**) Distribution (left) and morphology (right) of L2 ChCs in PL of an adult *Nkx2.1CreER:Ai14* mouse TM-induced at E17.5. Left: arrow indicates PL L2 ChCs. Right: dendrite (short arrow) soma (arrowhead) and axons (long arrow) of ChCs are indicated. Scale bar: 500 μm (left); 100 μm (right). AC: anterior cingulate cortex. (**b**) A neurolucida reconstruction of a single ChC sparsely labeled in a mouse with low dose TM induction (left; scale bar: 50 μm) and counts of total axon cartridges of reconstructed single ChCs (n=11, right). Plots indicate median, mean, quartiles and the range, the same in the following presentation. (**c**) Left: a schematic of labeling _BLA_PCs, _CC_PCs and _ST_PCs in PL by injecting 3 colors of retrograde CTB (Alexa 488, 594, 648) to 3 corresponding brain areas in the same mouse. Right: distribution patterns of 3 PC subsets in medial prefrontal cortex in single sections (100 μm thickness) and in overlay (1 mm thickness). Scale bar: 500 μm. (**d**) Average cortical depth of ChCs and 3 types of PCs in upper layers of the PL (cut off at 350 μm from pia) in the example section (_BLA_PC: 133.1±2.3 μm, n=63; _CC_PC: 230.3±6.1 μm, n=60; _ST_PC: 231.3±9.5 μm, n=52; ANOVA, p < 0.001). _BLA_PCs are located at similar laminar depth with ChCs (121.9±2.7 μm, n=69) but more superficial to _CC_PCs (p < 0.001, Mann-Whitney test). (**e**) Total number of cells that exhibit single or co-labeling of CTB, indicating specific or bifurcating axonal projections to injection sites. Of a total 6163 PCs counted, there were only 43 cells co-staining for BLA and CC projections. (**f**-**g**) Examples of synaptic responses from ChC to _BLA_PC and to _CC_PC. Upper panels: schematic of dual whole-cell patch recording of a ChC (red) and a _BLA_PC or _CC_PC labeled by CTB. Lower panels: representative traces from paired recordings in a _BLA_PC (green) or a _CC_PC (blue), showing unitary inhibitory postsynaptic currents (uIPSCs, averaged from 20–30 trials) evoked by paired action potentials (APs) in presynaptic ChCs. (**h**) Summaries of ChC-to-PC connection probability (numbers in graph indicate connected / tested pairs) and uIPSC magnitude (each circle represents individual connections).

To investigate synaptic connectivity between ChCs and _BLA_PCs or _CC_PCs, we performed paired wholecell patch recordings in L2/3 of PL in which ChCs expressed RFP and either _BLA_PCs or _CC_PCs were labeled with retrograde CTB-488 (Fig. 1f and Supplementary Fig. 3b). Strikingly, although _BLA_PCs and _CC_PCs had very similar morphological and intrinsic physiological features (Supplementary Fig. 4)^20^, L2 ChCs preferentially innervated _BLA_PCs over _CC_PCs indicated by both connection probability (87% vs. 17.5%, 20/23 pairs vs. 7/40 pairs; p < 0.01, Fisher exact test) and synaptic strength (ChC→_BLA_PCs: 36.0±4.0 pA, n=14; ChC→_CC_PCs: 11.1±3.4 pA, n=7; p < 0.001, Mann-Whitney test) (Fig. 1h). This highly selective ChC innervation of _BLA_PCs over _CC_PCs was not accounted by differences in their laminar location or distance from ChCs (Supplementary Fig. 5). Contrasting the suggestion from a previous study^9^, our results demonstrate remarkable selectivity of ChCs for PC subsets distinguished by projection target, though we cannot exclude the possibility that _CC_PCs might be more strongly controlled by another subset of ChCs.

### _BLA_PC-selective ChCs preferentially receive inputs from _CC_PCs

To examine local excitatory inputs to L2 ChCs, we recorded synaptic currents in ChCs following spikes evoked in either _BLA_PCs or _CC_PCs (Fig. 2a, b and Supplementary Fig. 6). Whereas 11.3% of _CC_PCs innervated ChCs (8 in 71 pairs, synaptic strength = 63.2±18.3 pA), only 1 _BLA_PC→ChC connection was observed in 60 tested pairs (p < 0.05, Fisher exact test) (Fig. 2c). This selective input from _CC_PCs over _BLA_PCs was even more striking considering that _BLA_PCs were located closer to ChCs than _CC_PCs (Fig. 1d).

**Figure 2.**
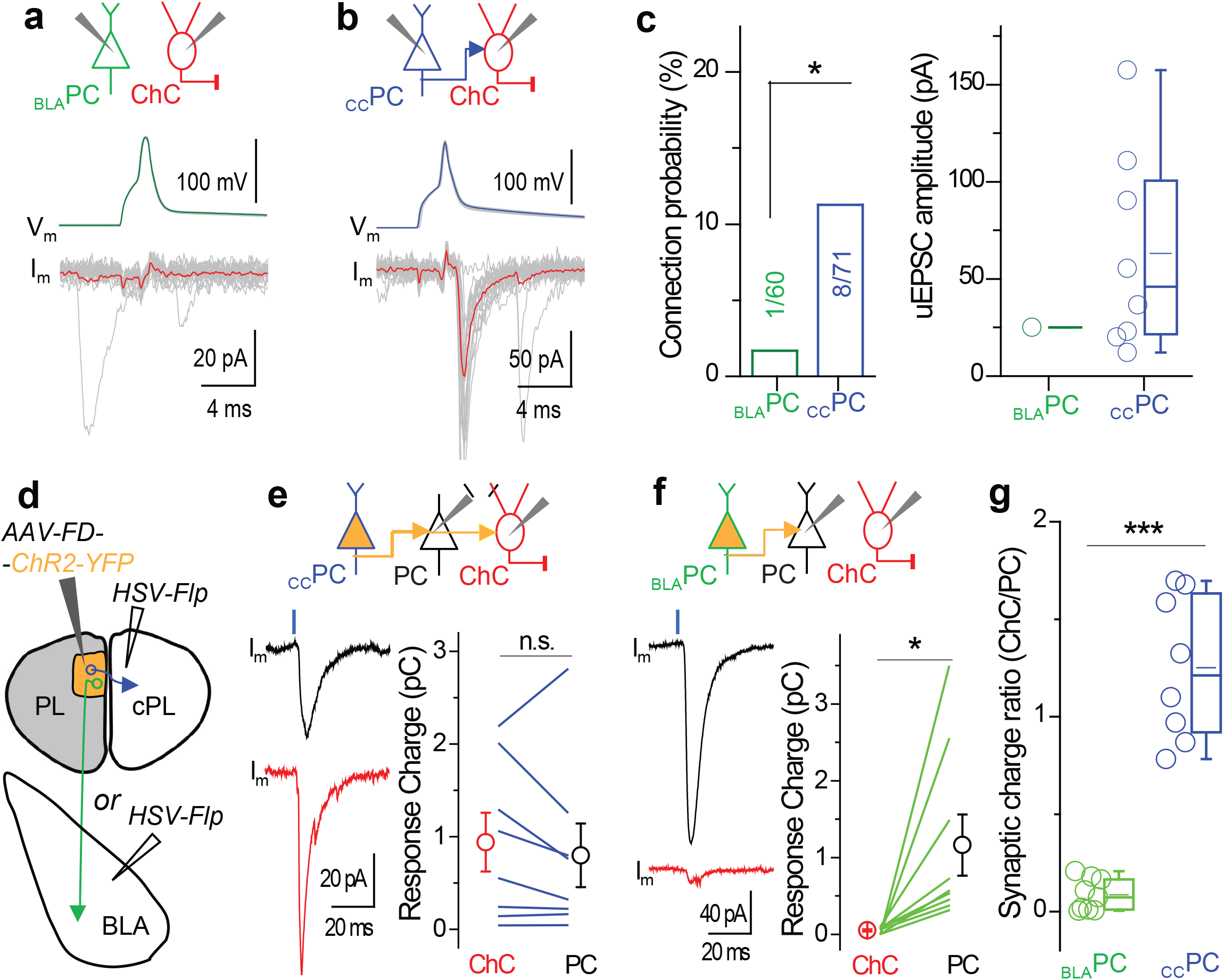
L2 ChCs receive strong input from _CC_PCs and weak input from _BLA_PCs. (**a**-**b**) Examples of synaptic responses in ChCs (red) following APs evoked in a _BLA_PC (**a**, green) and a _CC_PC (**b**, blue). Upper: schematic of dual recordings. Lower: representative traces from paired recordings in ChCs, averaged with thick traces from 20–30 trials. (**c**) Summaries of PC-to-ChC connection probability (numbers in bar graph indicate connected / tested pairs) and unitary excitatory postsynaptic current (uEPSC) magnitude, including results from experiments of loose-patch of presynaptic PCs (see Supplementary Fig. 6). (**d**) Schematic of dual viral delivery with retrograde *HSV-Flp* injection at contralateral PL (cPL) or ipsilateral BLA, respectively, followed by *AAV-FD-ChR2-YFP* injection in PL. (**e**) Upper panel: optical stimulation of _CC_PCs (blue) and whole cell recording of postsynaptic responses in adjacent pair of PC (black) and ChC (red). Lower left panels: example monosynaptic responses from a PC (black) and a nearby ChC (red) evoked by optical stimulation indicated by blue bars, averaged from 10 trials. Lower right panel: summary of synaptic response charges of paired neurons indicated by lines in _CC_PC expressing ChR2 (n=8), average values indicated by circles. (**f**) The same configuration as in **e** with optical stimulation of _BLA_PCs (green) (n=9). (**g**) Comparison of the ratio of synaptic response charge of ChCs over adjacent PCs following optical stimulation of the ChR2 axon from _BLA_PCs and _CC_PCs.

To assay inputs from broader populations of _CC_PCs and _BLA_PCs, we expressed channelrhodopsin-2 (ChR2) in each subset using a dual viral delivery strategy. A Flp-expressing retrograde herpes simplex virus (*HSV-Flp*) was first injected to either contralateral PL (cPL) or ipsilateral BLA; this was followed by the injection of a Flp-dependent ChR2-expressing adeno-associated virus (*AAV-FD-ChR2-YFP*) in PL to express ChR2 in _CC_PCs or _BLA_PCs, respectively (Fig. 2d and Supplementary Fig. 7; also see Methods). We then performed paired recordings of L2 ChCs and adjacent ChR2(−) PCs to measure the monosynaptic input from ChR2(+) PCs (see Methods). Optical stimulation of ChR2(+) _CC_PC axons evoked prominent monosynaptic responses in ChCs that were of similar strength to those in adjacent PCs (n=8 pairs; ChCs: 0.94±0.32 pC; PCs: 0.80±0.34 pC; p=0.35, Student’s paired t-test) (Fig. 2e). However, stimulation of _BLA_PCs evoked extremely weak synaptic responses in ChCs (n=9 pairs; ChCs: 0.05±0.01 pC; PCs: 1.16±0.40 pC; p=0.02, Student’s paired t-test) (Fig. 2f). We evaluated the strength of these inputs using the ratio of synaptic response charge in ChC vs PC for each pair (_CC_PC local input: 1.25±0.14; _BLA_PC local input: 0.09±0.03; p<0.001, Mann-Whitney test) (Fig. 2g). Thus L2 ChCs receive much stronger input from _CC_PCs than from _BLA_PCs, a recruitment specificity exactly opposite to their innervation specificity.

### ChCs and PVBCs form distinct microcircuit modules

As a comparison, we also assayed the connectivity pattern of PVBCs, which innervate the perisomatic region of PCs and also control PC output^22^. Using the *PV-Cre;Ai14* mice, we performed recordings in PL PVBCs and nearby CTB-labeled _CC_PCs or _BLA_PCs. L2/3 PVBCs innervated _BLA_PC and _CC_PC equally in both connection probability (37% vs. 34%, 10/27 pairs vs. 11/32 pairs; Pearson Chi-Square test: χ^2^=1.04, p=0.31) and synaptic strength (PVBC→_BLA_PCs: 34.5±15.7 pA, n=9; PVBC→_CC_PCs: 57.0±16.4 pA, n=7; p=0.46, Mann-Whitney test) (Fig. 3a). In the reverse direction, L2/3 PVBCs received equal inputs from _BLA_PC and _CC_PC in both connection probability (37% vs. 28%, 10/27 pairs vs. 9/32 pairs; Pearson Chi-Square test: χ^2^=1.84, p=0.17) and synaptic strength (_BLA_PC→PVBCs: 59.4±19.0 pA, n=8; _cc_PC→PVBC: 74.9±29.0 pA, n=7; p=0.95, Mann-Whitney test) (Fig. 3b). Thus, in contrast to PL ChCs (and hippocampal PVBCs^23^), PL PVBCs did not selectively connect with projection-defined PC subsets.

**Figure 3.**
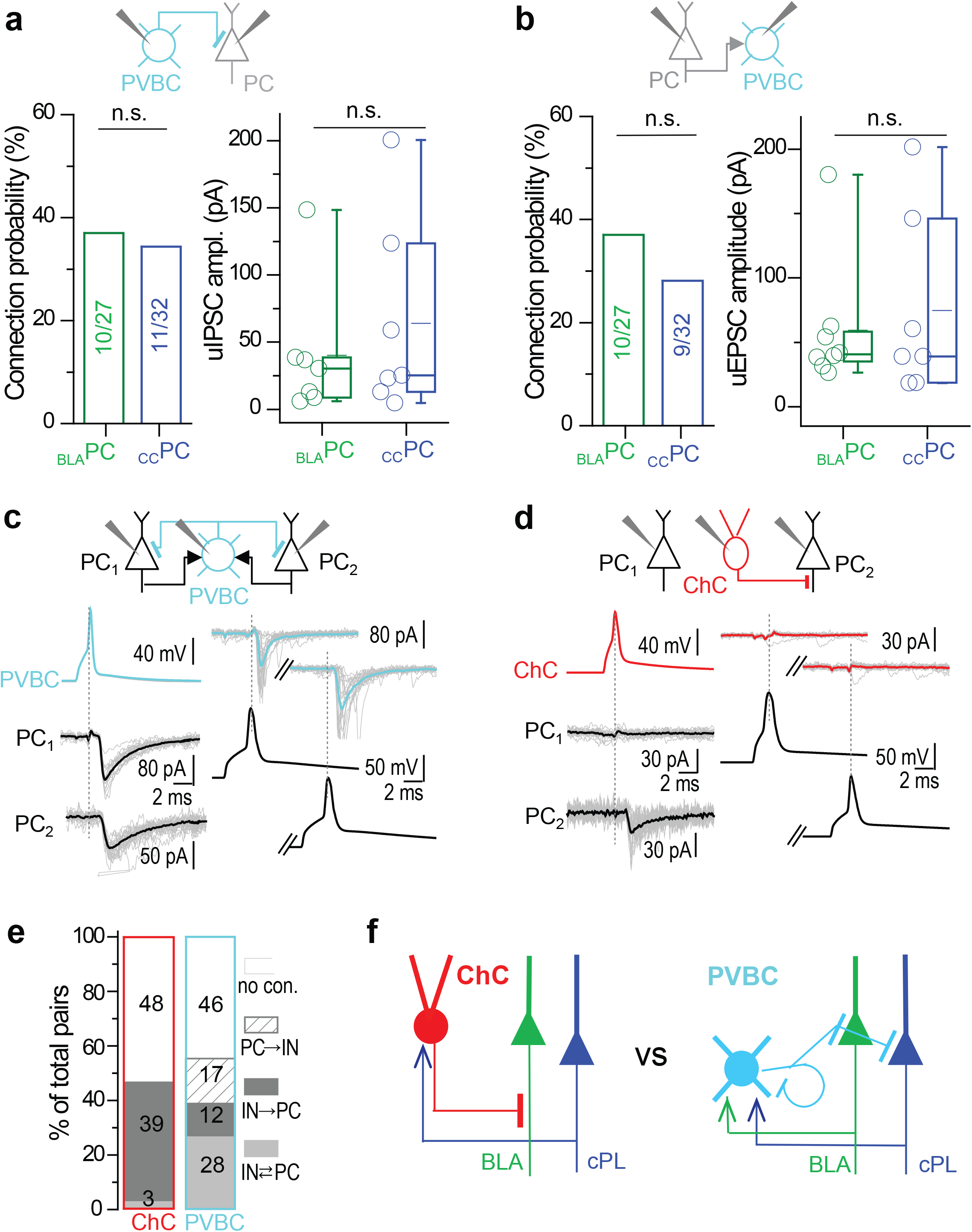
Distinct local circuit connectivity of PVBCs and ChCs in PL upper layers. (**a-b**) Summaries of PVBC-to-PC (**a**) and PC-to-PVBC (**b**) connection probability (numbers in graph indicate connected / tested pairs) and of the strength of uIPSC and uEPSC (each circle representing individual connections). Upper panels: schematic of dual whole-cell patch recording of a PVBC (cyan) and a PC (gray, _BLA_PC or _CC_PC) labeled by CTB. (**c**) Triple recording scheme and an example showing reciprocal connectivity in the PVBC microcircuit, with IPSC traces in both PCs evoked by PVBC spikes and EPSC traces in the PVBC evoked either PC spikes, averaged in thick trace from 15–20 trials. (**d**) Similar configuration as in **c** with ChC, instead of PVBC. (**e**) Summary of connectivity patterns in ChC or PVBC microcircuits in PL upper layer. Numbers in the bar graph indicate the number of pairs in each connection category. (**f**) Schematic of two distinct microcircuit modules mediated by ChC and PVBC, respectively.

To extend this comparison beyond connections with _BLA_PCs and _CC_PCs, we assayed ChC and PVBC connectivity with randomly selected PCs in PL upper layers (Fig. 3c-e). PVBCs formed extensive reciprocal connections with adjacent PCs (27.2%, 28 of 103 pairs), consistent with findings in other cortical areas^24,25^. In contrast, ChCs, while exerting extensive innervation of nearby PCs (synaptic kinetics shown in Supplementary Table 1), formed few reciprocal connections with these synaptic targets (3.3%, 3 of 90 pairs; p<0.001, Fisher exact test). This suggests a uni-directional connectivity pattern consistent with their sending output to _BLA_PCs and receiving input from _CC_PCs. Therefore, although both control PC output, ChCs and PVBCs form distinct inhibitory microcircuit modules (Fig. 3f).

### _BLA_PC-selective ChCs are preferentially recruited by the bilateral PL network

To systematically identify the local and long-range sources of input to L2 ChCs, we designed a genetic strategy that allowed trans-synaptic rabies tracing specifically from ChCs. To enable viral manipulation of ChCs, we generated a *Rosa26-loxpSTOPloxp-Flp* (*LSL-Flp*) mouse line that allowed conversion of transient *Nkx2.1*-driven *CreER* expression in progenitors of the medial ganglionic eminence to constitutive *Flp* recombinase expression in ChCs (Fig. 4a)^26^. A modified trans-synaptic tracing strategy involving 2 AAV helpers and a glycoprotein-deleted (dG) rabies virus was used to reveal the overall pattern of local and long-range monosynaptic inputs to L2 ChCs in PL (Fig. 4b, also see Methods). We used GAD67 immunostaining to distinguish GABAergic vs glutamatergic (i.e. GAD67(−)) neurons labeled by EnvA-dG-GFP. Within the cortex, presynaptic GABAergic neurons (including those positive for PV and vasoactive intestinal peptide) were distributed across cortical layers with slight enrichment in L1 (GAD67(+): L1, 5.9±1.4%; L2, 4.5±0.8%; L3, 4.6±0.3%; L5/6, 1.9±0.5 % of total local inputs; Fig. 4c-e and Supplementary Fig. 8). On the other hand, presynaptic glutamatergic neurons (GAD67(−): L1, 4.7±1.2%; L2, 8.2±1.1%; L3, 40.6±4.6%; L5/6, 30.7±4.2% of total local inputs) were sparse in L2 but were more enriched in L3 and L5/6 (Fig. 4d, e), suggesting that L2 ChCs received much fewer excitatory inputs from adjacent PCs in the same layer than PCs in more distant layers, consistent with the paired recording results (Fig. 1, 2).

**Figure 4.**
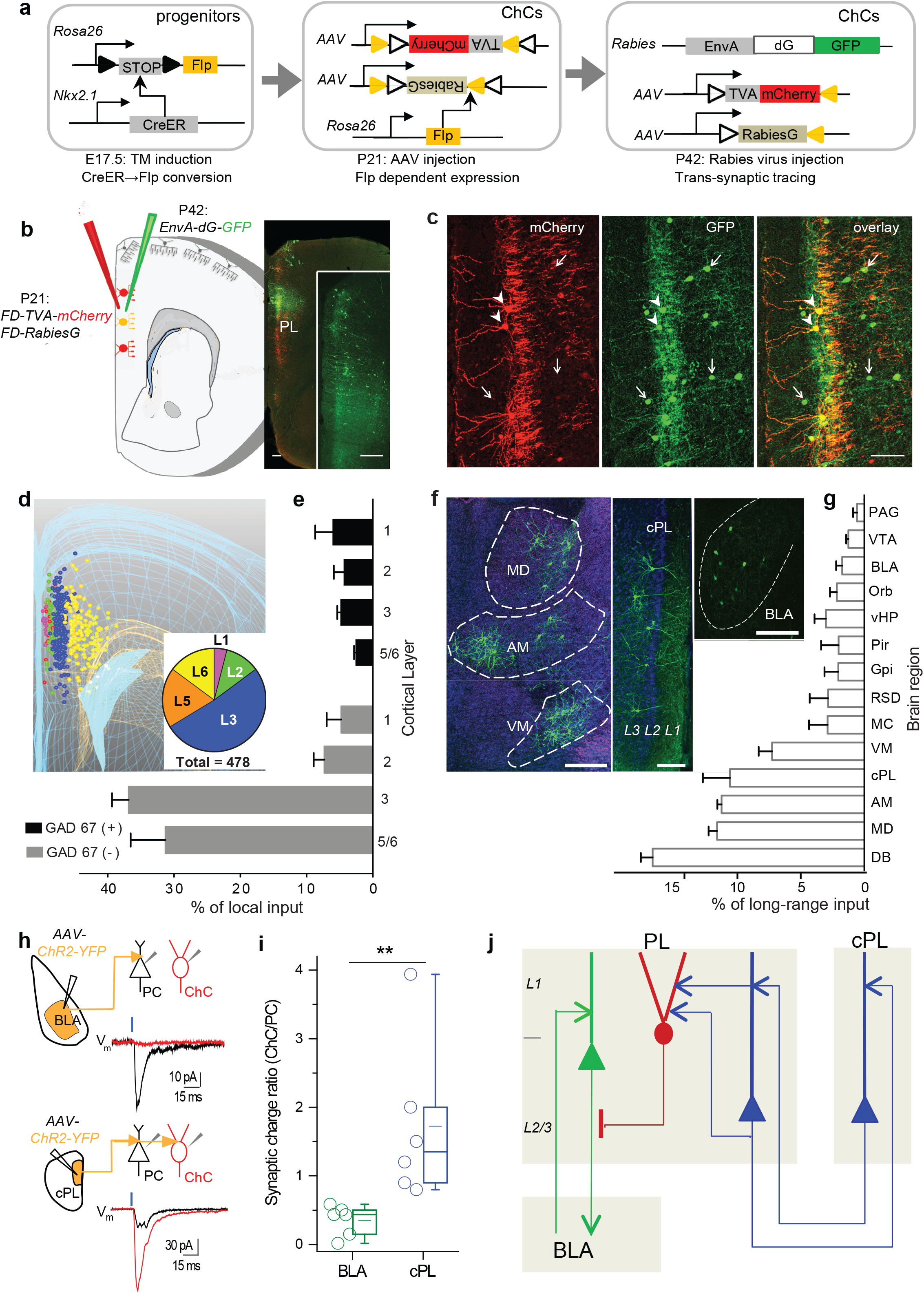
Systematic tracing of local and long-range inputs reveal that L2 ChCs are preferentially recruited by bilateral _CC_PC input as opposed to BLA input. (**a**) Scheme of converting transient *CreER* activity in Nkx2.1(+) progenitors to permanent *Flp* activity in ChCs that enables AAV and rabies viral targeting in mature cortex. (**b**) Left: Schematic of trans-synaptic rabies tracing specifically from PL L2 ChCs. Right: overview of PL region triple-infected with *AAV-FD-TVA-mCherry, AAV-FD-RabiesG*, and *rabies-EnvA-dG-GFP*. Inset: overview of PL region with GFP expression. Scale bar: 200 μm. (**c**) Example of retrograde trans-synaptic tracing from L2 ChCs in PL. Triple-infected ChC starter cells coexpress mCherry and GFP (arrowheads); some of their pre-synaptic cells incorporated GFP through *rabies-GFP* (arrows). Scale bar: 100μm. (**d**) A 3D stereological rendering/reconstruction of total local inputs to PL L2 ChCs from a single tracing experiment. The laminar distributions of pre-synaptic cells are colored and their ratios presented as a pie chart. (**e**) Laminar source of local input to L2 ChCs separated by GABAergic or glutamatergic cells (n=5 mice, see Supplementary Fig. 8). (**f**) Examples of long-range input sources to PL L2 ChCs, indicated by GFP labeling in several thalamic nuclei, cPL and BLA. Scale bar: 100 μm. (**g**) Distribution of long-range inputs to PL L2 ChCs (n=5 mice) measured as percentage of the total number of presynaptic cells detected brain wide. (DB: diagonal band; MD, AM, VM: mediodorsal, anteromedial, ventromedial thalamic nuclei; MC: motor cortex; RSD: retrosplenial area dorsal part; Gpi: globus pallidus internal segment; Pir: piriform area; vHP: ventral hippocampus; Orb: orbitofrontal cortex; VTA: ventral tegmental area; PAG: periaqueductal gray.) (**h**) Examples of synaptic responses in pairs of adjacent ChCs (red) and PCs (black) evoked by optical stimulation (blue bars) of ChR2-expressing BLA (upper panels) or cPL (lower panels) input axons. Left schematics depict stimulation and recording configurations following *AAV-ChR2* infection of BLA or cPL. Synaptic responses were averaged from 10 trials. (**i**) Summary of the ratio of synaptic response charge of ChCs over adjacent PCs following optical stimulation of the ChR2 axon from BLA (n=6 pairs) or cPL (n=6 pairs). (**j**) A schematic model depicting that L2 ChCs in the PL mediate directional inhibition in local circuits from _CC_PCs to _BL_APCs and in global networks from the bilateral _CC_PC reciprocal network to the PL-BLA reciprocal network.

Major sources of long-range input to L2 ChCs (Fig. 4f, g) included the diagonal band of the basal forebrain (16.3±0.9%, including cholinergic input, Supplementary Fig. 9), the mediodorsal (11.7±0.7%), anteromedial (11.3±0.3%) and ventromedial (7.3±1.0%) thalamic nuclei, and contralateral PL (cPL; 10.6±2.1%). Although the BLA prominently projects to PL as part of a PL-BLA reciprocal network ^20^, it was a relatively minor source of long-range inputs to L2 ChCs in the PL (BLA input: 1.8±0.5%). To validate the physiological connections and possible selectivity of long-range inputs, we employed optogenetic measurements to compare the synaptic strength from cPL and BLA to L2 ChCs. Following *AAV-ChR2-YFP* injection into BLA or cPL in mice in which ChCs expressed RFP, we performed paired recordings of L2 ChCs and adjacent PCs in the PL upon light activation of ChR2(+) BLA or cPL axons, respectively (Fig. 4h). We evaluated the strength of these inputs using the ratio of synaptic response charge in ChC vs PC for each pair. Whereas cPL provided strong input to ChCs (1.72±0.52, n=6 pairs), BLA sent weak input (0.35±0.10, n= 6 pairs; p<0.01, Mann-Whitney test) (Fig. 4i). Therefore, L2 ChCs in PL not only receive stronger input from local _CC_PCs than from _BLA_PCs, but also receive stronger input from _CC_PCs in cPL than from projection neurons in the BLA. On the other hand, BLA neurons project stronger input to _BLA_PCs than _CC_PCs in the superficial layers, and contralateral _CC_PCs provide similar input to _BLA_PCs and _CC_PCs^20^. Taken together, these data suggest that the ChCs are preferentially recruited by the reciprocal network comprised of callosally-projecting _CC_PCs from the two hemispheres (PL-cPL network), compared to the reciprocal PL-BLA network (Fig. 4j).

### L2 ChCs suppress PC firing in freely behaving mice

To examine the physiological impact of ChCs on PCs *in vivo*, we combined optogenetic manipulation of ChCs with single unit recording of PCs in freely behaving mice. We virally expressed ChR2 in L2 ChCs by injecting *AAV-FD-ChR2-YFP* into the PL of mice expressing *Flp* in ChCs (Fig. 4a). Of total virally labeled cells, 88.6±5.6 % cells were ChCs; in layer 2/3 specifically 94.8±3.6% of cells were ChCs (n=5 mice, Supplementary Fig. 10). We implanted optrodes targeting the PL and field electrodes targeting the ipsilateral BLA (Fig. 5a, b, also see Methods). In the PL, 79 well-isolated single units were recorded in awake, quietly resting mice (n=3) while they freely explored a small box. Following brief (5 ms, 5 mW) pulses of blue light delivered at 1 Hz, 3 of the 79 units showed robust short latency (3–4 ms) excitatory responses (Fig. 5c-e), suggesting that they were likely to be ChR2-expressing ChCs directly activated by light. A larger number of units (13/79, 16.5%) were inhibited at typically longer latencies (ranging from 2-13 ms) (Fig. 5c, f-g), suggesting that these neurons received monosynaptic inhibition from ChR2-expressing ChCs. Trial-by-trial analyses showed that such inhibition was independent of the baseline firing state of the unit (Supplementary Fig. 11). Interestingly, 3 of these inhibited units had latencies (2–3 ms) even shorter than those seen in the 3 putative ChCs. These short latency inhibitory responses may have resulted from direct activation of ChC boutons along the AIS of these units, consistent with a fast and powerful inhibition at the spike initiation site. No significant short-latency firing rate responses to light were observed in 65 neurons recorded from control animals (n=3 mice) expressing eYFP in ChCs.

**Figure 5.**
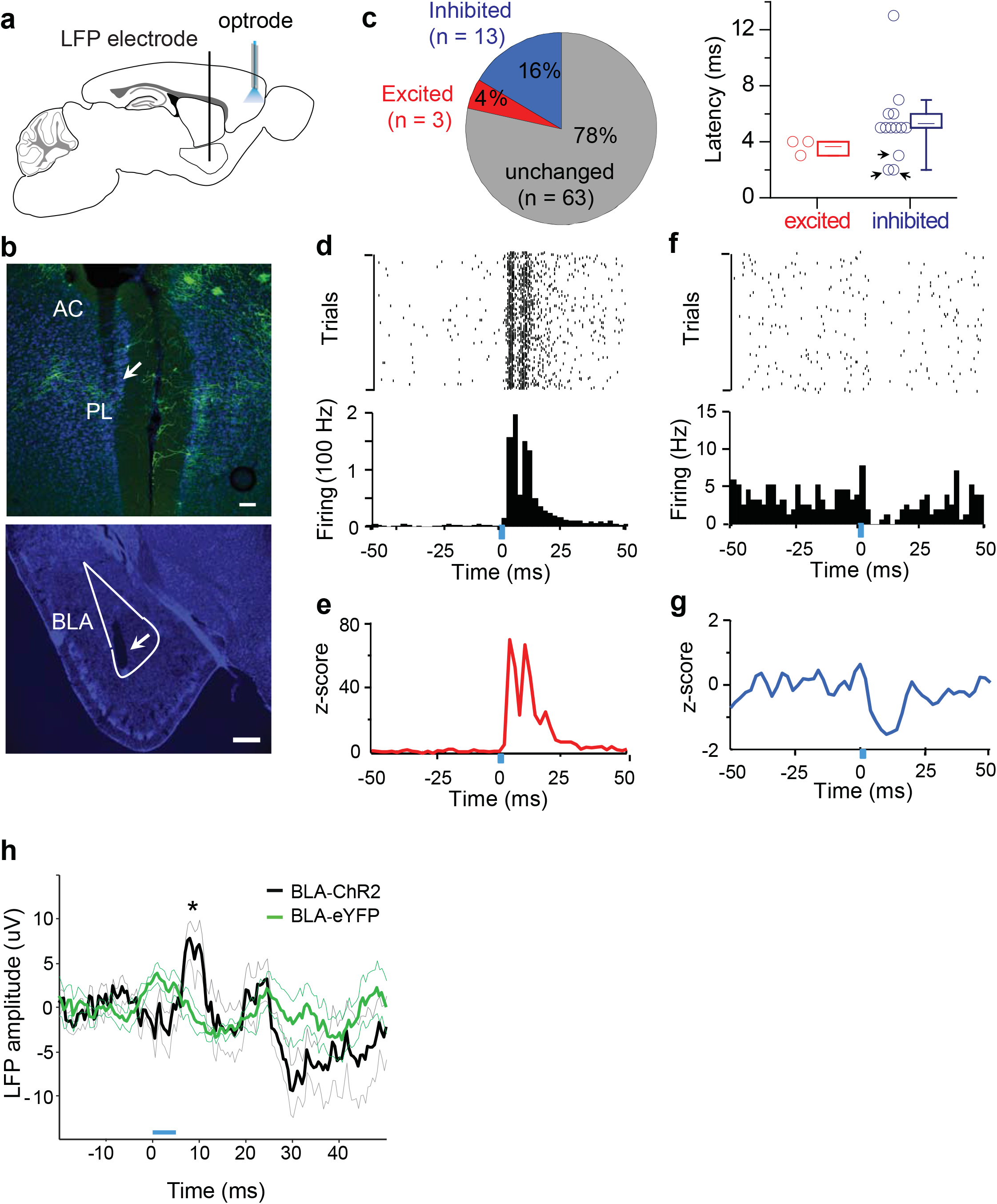
Optogenetic activation of L2 ChCs in PL inhibits PL firing, including _BLA_PC firing, in freely behaving mice. (**a**) Schematic of optrode stimulation and recording in the upper layer of PL with simultaneous local field potential (LFP) monitoring in ipsilateral BLA. (**b**) A coronal section from a *NkxCreER;LSL-Flp* mouse brain with bilateral infection of *AAV-FD-ChR2-YFP* in the PL. Arrows indicate the optrode track through the frontal cortex (upper panel) and the electrolytic lesion of LFP electrode in the BLA (lower panel). Green, ChR2; blue, DAPI; scale, 100 μm; AC, anterior cingulate. (**c**) Percentages (left) and latencies (right) of statistically significant excited or inhibited PL single units. Significance determined by bootstrapping (see Methods). Note the three inhibited units with very short latency (≤ 3ms, indicated by arrows) (see text). (**d**) Spike raster (top) and peri-stimulus time histogram (PSTH, bottom) for a light-excited unit that increased firing within 15ms of light onset (blue bar). Light pulse duration, 5ms; frequency, 1Hz. (**e**) Average PSTH for the 3 short-latency light-excited units. (**f-g**) Same as in **d-e** for short-latency light-inhibited units. (**h**) Evoked potential in BLA during the optical stimulation of PL. Blue rectangle indicates light pulse. Average evoked potential 5–10ms following light pulse was significantly different between control and the ChR2 group (ChR2: 5.58±3.56 μV, n=6 sessions from 3 mice; eYFP: −0.38±3.57 μV, n=5 sessions from 3 mice; p<0.05, two-sample t-test).

To assess whether PCs suppressed by ChCs included _BLA_PCs, we first examined the impact of ChC activation on BLA local field potential (LFPs). Our analysis revealed a fast and robust stimulation-evoked positive-going (likely inhibitory) evoked response in the BLA LFP 5–10ms following light stimulation in PL (Fig. 5h), suggesting that ChC activation suppressed the activity of the PL projection to the BLA. Next, we examined the recordings for evidence of connectivity between the recorded PL single units and the BLA, by examining the phase-locking of PL single unit spikes to BLA 3–6 Hz LFP oscillations^27,28^. Overall, 13/79 (16.5%) PL units were significantly phase-locked to the BLA LFP (p<0.05, Rayleigh’s test of circular uniformity, see Methods). Importantly, the fraction of light-inhibited units that was phase-locked to the BLA LFP (4/13 or 31%) was significantly higher than that of units that did not change firing upon ChC activation (9/63 or 14%; p<0.05, χ_2_ = 4.84, Pearson two-sample Chi-square test) (Supplementary Fig. 12a, b). Light-inhibited units (8/13 or 62%) were also more likely than other units (29/58 or 50%) to be more strongly phase-locked to the BLA LFP of the future, suggesting a net PL-to-BLA directionality specifically in the inhibited units (Supplementary Fig. 12c, d)^29^. Together, these results corroborate the *in vitro* experiments, suggesting that ChCs preferentially inhibit _BLA_PCs *in vivo.*

## DISCUSSION

A fundamental question in understanding the functional organization of cortical circuits is whether diverse GABAergic neurons mediate more or less the same non-selective, “blanket” inhibition or contribute to specialized connectivity motifs that shape PC subnetworks underlying specific forms of circuit operations and information processing^4,30,31^. One set of studies suggested a general lack of target selection for neocortical interneurons^32,33^, but these studies mostly did not distinguish bona-fide interneuron types nor PC subsets. Although certain interneurons may indeed mediate non-selective inhibition in certain circuit contexts, e.g. neurogliaform cells^34,35^, several studies have reported selectivity of GABAergic neurons for PC subpopulations in cortex and hippocampus^23,36–38^. In particular, despite the striking subcellular selectivity of ChC innervation^6,7,9^, their circuit connectivity pattern is poorly understood^9,11^. Here, by capturing a subset of a bona-fide interneuron type and projection-defined PCs, we demonstrate exquisite specificity in the directional innervation as well as recruitment of ChCs not only in local circuits but also in global networks. This directional ChC module may promote physiological segregation of intermingled _CC_PC and _BLA_PC ensembles. As _CC_PC and BLAPC are each embedded in distinct larger scale networks, this might provide a cellular basis for hierarchical control of one brain network (the PL-cPL network) over another (PL-BLA network). These results suggest that the specialization of interneuron subpopulations in the inhibitory control of discrete PC ensembles might be a key principle of cortical organization. These ensembles might then be combined in order to construct hierarchical or parallel information processing streams in global networks. Defining the features and degrees of inherent specificity of such connectivity templates will thus provide biological ground truth for building models of cortical computation and information processing.

A key prerequisite to discovering the specificity of neural connectivity is the identification of appropriate neuronal subpopulations or subtypes – basic building blocks of circuit motifs and network scaffolds^39^. While multiple major classes or populations of cortical GABAergic neurons have been recognized, the specific subpopulations that constitute functional circuit modules remain largely unknown. In the hippocampus, a recent set of studies demonstrates that PVBCs consist of subpopulations with distinct embryonic birth date, input connectivity, and output target neurons, each of which play distinct roles in network level plasticity and learning^40,41^. In this context, it should be noted that cortical ChCs are generated from *Nkx2.1* progenitors during late gestation mainly between E15.5 and E18.5^21^, and our current results on their wiring specificity are derived from a subset of L2 ChCs born at E17.5 or later. It is possible that earlier-born ChCs might exhibit different selectivity for PCs, such as _CC_PCs and/or other PC subsets in PL. As E15.5 *Nkx2.1* progenitors generate a large proportion of non-ChCs, a more refined genetic tool that targets earlier-born ChCs will facilitate examining this intriguing possibility.

The extensive control of PC firing by PV- and CCK-positive basket interneurons, both innervating the perisomatic region^42^, raised intriguing questions about the role of ChCs that target the AIS. Here we demonstrate that, beyond their differences in subcellular selectivity, ChCs and PVBCs differ substantially in their local connectivity and therefore represent distinct microcircuit modules. While PVBC-PC connectivity is extensively reciprocal and largely non-selective, connectivity between ChCs and PCs is directional and highly selective. While the multipolar dendrites of L2/3 PVBCs receive dense inputs from local PCs, the predominant apical layer 1 dendrites of ChCs receive rather sparse local excitatory inputs but more extensive inputs from other cortical layers and diverse long-range sources. ChCs and PVBCs further differ in their *in vivo* spike timing during brain state transitions and coupling to network oscillations^43^. Together, these results suggest that PVBCs are well suited to regulate the balance, gain and network oscillation of relatively broad PC populations^22,42^. Instead, ChCs may mediate the dynamic segregation and hierarchical interaction of select PC ensembles, thereby routing information flow through local circuits and global networks, especially in response to more distant and long-range inputs.

Importantly, we provide the first compelling evidence that ChCs inhibit PC firing *in vivo.* Our results do not exclude the possibility that ChCs might also exert excitatory effects under certain network states (e.g. the downstate) when the PC AIS is substantially more hyperpolarized from the chloride equilibrium potential^14^. Future studies that monitor and manipulate ChCs in behavioral tasks that engage different brain states will further clarify the cellular impact of ChCs in orchestrating dynamic PC ensembles and circuit operations^44^.

In associative fear learning, the activity of PVBCs in PL is modulated by conditioned stimulus and contributes to the synchronization of PC firing (including _BLA_PC) that drives fear expression^45,46^. Given their selective and directional inhibition of _BLA_PC and likely the PL-BLA network, ChCs in contrast to PVBCs, may suppress fear expression according to “upstream signals”. In this context, our finding that ChCs receive major inputs from the bilateral _CC_PC network and the mediodorsal thalamus (MD) is notable. As a high-order thalamic nucleus, MD integrates inputs from orbitofrontal cortex, medial frontal cortex, BLA, and basal ganglia, projects to prefrontal cortex^17,18,47^, and has been implicated in working memory and cognitive flexibility^48^. It is thus possible that an inhibitory control of the _BLA_PC-BLA network by the MD-CCPC network through L2 ChCs might contribute to cognitive and flexible regulation of fear expression.

## ACKNOWLEGEMENT

We thank Selin Schamiloglu for help with retrograde labeling of PL pyramidal neurons, Bo Li, Adam Kepecs and Gyuri Buzsaki for comments on the manuscript. This work was supported in part by NIH R01 MH094705-05 and CSHL Robertson Neuroscience Fund to Z.J.H., and by NIH R01 MH081968 and the Hope for Depression Research Foundation to J.A.G. J.T. was supported by NRSA F30 Medical Scientist Predoctoral Fellowships. J.L. was supported by a NARSAD Postdoctoral Fellowship. N.P.C. was supported by the National Science Foundation.

## Supplementary Materials

### Methods

#### Experimental animals

In order to genetically label and manipulate chandelier cells, we crossed *Nkx2.1-CreER* mice (The Jackson Laboratory stock 014552) with either *Rosa26-lox-stop-lox-TdTomato (Ai14)* reporter (The Jackson Laboratory stock 007905) or in house derived *Rosa26-lox-stop-lox-Flp (LSL-Flp)* mice^1^. To properly identify embryonic day 17.5 (E17.5) for tamoxifen (TM) inductions, Swiss Webster females (Taconic) were housed with *Nkx2.1CreER;Ai14 (het/homo)* males overnight and females were checked for vaginal plug by 8–9 am the following morning. Positive plug identification was timed at E0.5. To genetically label pavalbumin positive basket cells (PVBCs), we crossed *PV-Cre* mice (The Jackson Laboratory stock 008069) with *Ai14* reporter. The ages of animals used are e indicated in the different experiments stated below. Both male and female mice were employed without distinction in all the experiments. All experiments were conducted in accordance with the Institutional Animals Care and Use Committee of Cold Spring Harbor Laboratory.

#### TM induction

TM was dissolved in corn oil (20 mg/ml) overnight, at room temperature under constant stirring. Stocks were stored as individual aliquots at 4°C degrees for no more than one month. Following light isoflurane anesthesia pregnant females were given oral gavage administration of TM (dose: 3 mg / 30 g of body weight) at gestational day E17.5 for dense revealing of ChCs. In rare instances, TM induction lead to dystocia in pregnant females and emergency caesarian sections were performed. Pups retrieved following caesarians were housed with Swiss Webster foster mothers until weaning age. For the experiment of sparse labeling of ChCs for single cell reconstruction, the low dose of TM (0.1 mg / 30 g of body weight) was used.

#### Viral Constructs

*HSV-Elf1a-Flp* was purchased from Rachael Neve, Viral Gene Transfer Core, MIT; *AAV-Elf1a-FD-ChR2-YFP* was a gift from Deisseroth lab, Stanford University. *AAV-CAG-ChR2-YFP* was purchased from UNC vector core, Chapel Hill, North Carolina. *Ef1a-FD-TVA-mcherry* (TVA: avian glycoprotein EnvA receptor), *Ef1a-FD-RabiesG* (RabiesG: rabies glycoprotein) cassettes were assembled and cloned using standard molecular cloning protocols with restriction enzymes from New England Biolabs. (1) TVA-mcherry (pAAV-EF1a-FLEX-TVA-mCherry was a gift from Naoshige Uchida (Addgene plasmid # 38044)); (2) RabiesG (pAAV-CA-FLEX-RG was a gift from Naoshige Uchida (Addgene plasmid # 38043)). Each assembled cassette was subcloned into the *AAV-Ef1a-FD-YFP-WPRE* (a gift from Deisseroth lab, Stanford University)^2^, using NheI and AscI cloning sites. All constructs were sequenced to ensure their fidelity and proper reversed orientation of given inserts, and packed into AAV8 viral vectors with titers ranging from 1.0−2.4 × 10^12^ pfu from UNC Vector Core (Chapel Hill, North Carolina). A pseudotyped rabies virus expressing the avian glycoprotein EnvA (*EnvA-dG-GFP*, 4.3 × 10^8^ pfu) was purchased from Salk GT3 Vector Core (La Jolla, California).

#### Surgical Procedures of stereotaxic injection

Animals were anesthetized by an intraperitoneal injection of ketamine/xylazine (dose: 100 mg/kg ketamine, 10 mg/kg xylazine in saline). Mice were mounted in a stereotaxic headframe (Kopf Instruments Model 940 series). Bregma coordinates were identified for 4 brain areas: PL and contralateral PL(cPL) (antero-posterior, A/P: 2.0 mm; medio-lateral, M/L: 0.2−0.3 mm; Dorso-ventral, D/V: 1.5 mm depth from the pial surface), BLA (A/P: −1.6mm, M/L: 3.0-3.25 mm; D/V: 3.75 mm), dorsomedial striatum (A/P: 1.0 mm; M/L: 1.2 mm; D/V: 2.25). An incision was made over the scalp, a small burr hole was made into the skull and brain surface was exposed. A pulled glass pipette tip of 20–30 μm containing virus or tracers was lowered into the brain. Pulses were delivered using a Picospritzer (General Valve Corp) at a rate of 30 nl/minute; the pipette was left in the brain for 5–10 minutes to prevent backflow^3^. Following the injection, the pipette was withdrawn, the incision was closed with tissue glue, and animals recovered.

#### Retrograde tracing of PC subtypes

For anatomical characterization of PC subsets in PL region, the retrograde neuronal tracing of cholera toxin B subunit (CTB) was used. Three colors of CTB (Alexa Fluor-488, 594 and 647) (Life Technologies, 0.3 μl, 2% in PBS) were injected into BLA, dorsomedial striatum, and cPL in the same mouse, respectively. The laminar distribution and co-staining analysis performed as described below.

For physiological paired recordings, CTB-488 (0.3 μl, 2% in PBS) was injected into either BLA or cPL to label _BLA_PC or _CC_PC in the PL upper layer. We post-hoc verified the proper placement of injection site for physiological experiments. For all CTB experiments animals were either perfused or prepared for slice physiology 5–10 days post injection.

#### Retrograde rabies tracing

##### Virus injection

A modified rabies virus strategy was used involving the AAV helper virus and a glycoprotein-deleted (dG) rabies virus to trace monosynaptic inputs to ChCs (Fig. 4a, b). In *Nkx2.1CreER;LSL-Flp* mice (TM induction at E17.5), ChCs express Flp in mature ages. At P21, 2 AAV viruses mixture of *FD-TVA-mcherry* and *FD-RabiesG* (1:1, 0.3 μl) was injected into unilaterally PL. Flp-expressing ChCs activated the AAV vectors and expressed TVA and RabiesG. Three weeks following AAV injections, 0.3 μl of *EnvA-dG-GFP* rabies virus was injected into the same PL coordinates so that only the TVA-containing ChCs were infected. The modified rabies encodes GFP instead of its native glycoprotein. Cells infected with rabies virus were labeled with mCherry and GFP, allowing for their easy identification. These “starter cells” also expressed RabiesG from the AAV vector, and allowed monosynaptic retrograde spread of dG rabies virus. Presynaptic cells expressing GFP from the rabies virus were easily distinguished from the mCherry-labeled starter cells. Animals recovered for 1 week before perfusion to allow sufficient local and long range labeling with pseudotyped rabies virus. In some cases, unpseudotyped rabies injections were used for retrograde labeling of PC (_BLA_PC or _CC_PC) subtypes, 0.5μl of *dG-GFP* rabies virus was injected unilaterally into cPL or BLA.

##### Histology

Seven days following rabies infection, animals were perfused with 4% PFA in PBS. Brains were removed and postfixed overnight using the same fixative. Coronal brain slices were sectioned at a 100 μm thickness using a vibratome. For histological analysis of local microcircuiry using GABAergic makers, and analysis of AIS GABA boutons, slices either through the injection site (in the case of rabies input mapping) or PL (in the case of AIS analysis) were sectioned at 20μm. Sections were blocked with 10% normal goat serum in 0.5% Triton in PBS and then incubated overnight with combinations of the following primary antibodies diluted in block solution: rabbit polyclonal RFP (1:1000, Rockland) or chicken polyclonal anti-GFP (1:1000, Aves) for fluorophore preservation of mcherry starter cells and GFP rabies virus expression, mouse monoclonal anti-parvalbumin (1:1000 Sigma), rabbit polyclonal phospho-IkappaB (1:300, Cell Signaling), rabbit polyclonal VIP (1:250, Immunostar), rabbit polyclonal somatostatin-14 (1:500, Peninsula), mouse monoclonal GAD-67 (1:500, EMD Millipore), mouse monoclonal VGAT (1:500, synaptic systems), goat polyclonal ChAT (1:500, EMD Millipore), and rabbit polyclonal NeuN (1:1000, Abcam).

For immunostaining with GAD67, in order to see somatic labeling for GABA positive inputs, no detergent was used but detergent was added in samples were GAD67 boutons were analyzed at the AIS. Sections were incubated with appropriate Alexa fluor dye-conjugated IgG secondary antibodies (1:500, Molecular Probes). In some instances, for identification of distal brain structures such as inputs from particular nuclei of the thalamus and basal forebrain, sections were incubated with Neurotrace Fluorescent Nissl Stain in secondary antibody (1:300, Molecular Probes). Sections were washed and mounted with Fluoromount-G (Southern Biotech).

#### Image Acquisition and Analysis

##### Input tracing

Images were taken by confocal microscopy (Zeiss LSM 780). All images were processed using Fiji^4^. For local input analysis, serial sections through the PL cortex were acquired. In some instances images were spatial registered using BUnwarp J Fiji plugin^5^. Individual images with NISSL signal were manually overlaid in Photoshop with representative atlas images (Paxinos and Watson Mouse Brain in Stereotaxic Coordinates, 3rd edition). For 3D model reconstructions and visualization, image stacks were uploaded and manually traced with open source software FreeD ^6^. Cortical depth was gauged based on arbitrary assignment of cortical depths (layer 1 = 100 μm, layer 2 = 100 − 200 μm, layer 3 = 200 − 400 μm, layer 5/6 = great than 400 μm depth).

##### Single ChC reconstruction

For single cell reconstruction in sparse labeling of ChCs, individual cell morphology was traced using Neurolucida software packages (Microbrightfield). Bouton analysis at the AIS of PCs was done with 63x oil immersion lens at a zoom factor of 2.1 and followed previous described protocols^7–9^. Briefly, AIS were identified by rabies-GFP label in an axonal process co-localized with phosphor-IkappB that was connected to a pyramidal soma. Inhibitory boutons were defined as 0.5-1μm varicosities within more than one 0.3μm thick imaging plain and positioned 1 μm or less from an AIS.

#### In vitro Electrophysiology

##### Slice preparation

We used *Nkx2.1-CreER;Ai14* or *PV-Cre;Ai14* mice to investigate the circuit organization of ChC or PVBC network in the PL. Mice (>P30) were anesthetized with isoflurane before decapitation.

The dissected brain was rapidly immersed in ice-cold, oxygenated, artificial cerebrospinal fluid (section ACSF: 110 mM choline-Cl, 2.5 mM KCl, 4mM MgSO4, 1mM CaCl2, 1.25 mM NaH2PO4, 26mM NaHCO3, 11mM D-glucose, 10 mM Na ascorbate, 3.1 Na pyruvate, pH 7.35, 300 mOsm) for 1 min. Coronal prefrontal cortical slices were sectioned at 300 μm thickness using a vibratome (HM 650 V; Microm) at 1-2 °C and incubated with oxygenated ACSF (working ACSF; 124mM NaCl, 2.5 mM KCl, 2 mM MgSO4, 2 mM CaCl2, 1.25 mM NaH2PO4, 26 mM NaHCO3, 11 mM D-glucose, pH 7.35, 300mOsm) at 34 °C for 30 min, and then transferred to ACSF at room temperature (25 °C) for >30 min before use. Whole cell patch recordings were directed to the medial part of frontal cortex (including PL), the morphology of subcortical whiter matter and corpus callosum as primary landmarks according to the atlas (Paxinos and Watson Mouse Brain in Stereotaxic Coordinates, 3rd edition).

##### Electrophysiological recordings

Patch pipettes were pulled from borosilicate glass capillaries with filament (1.2 mm outer diameter and 0.69 inner diameter; Warner Instruments) with a resistance of 3–6 MΩ. The pipette recording solution consisted of 110 mM potassium gluconate, 30 mM KCl, 10 mM sodium phosphocreatine, 10 mM Hepes, 4 mM ATP·Mg, 0.3 mM GTP, and 0.3 mM EGTA (pH 7.3 adjusted with KOH, 290 mOsm). Dual or triple whole cell recordings from RFP labeled cells at the L1/2 border and CTB-488 labeled PCs in layer 2/3 were made with Axopatch 700B amplifiers (Molecular Devices, Union City, CA) using an upright microscope (Olympus, Bx51) equipped with infrared-differential interference contrast optics (IR-DIC) and fluorescence excitation source. In some experiments, PCs, identified by their triangle soma and thick primary dendrite, were blindly selected within 100 μm distance to RFP positive interneuron without retrograde tracer. Both IR-DIC and fluorescence images were captured with a digital camera (Microfire, Optronics, CA). All recordings were performed at 33–34 °C with the chamber perfused with oxygenated working ACSF.

Synaptic connection was detected as similar as in^10^. Synaptic responses were evoked by presynaptic action potentials (APs) through soma-injected current squares (1.5–3 ms, 1–2.8 nA). In some experiments, we evoked APs in PCs by loose patch stimulation. The loose patch was achieved by the tight touch (> 100 MΩ resistance) to the targeted PCs through the same pipette for the whole cell patch recordings. The stimulation ranged from 0.1–1 V in 200 μs, 0.1Hz. The intensity used was determined by the persistent spikes after the stimulation because of the immediate spike masked by the artifact produced by the stimulation. Recordings were made with two MultiClamp 700B amplifiers (Molecular Devices). The membrane potential was maintained at −75mV in the voltage clamping mode and zero holding current in the current clamping mode, without the correction of junction potential. The postsynaptic neurons were held at −75 mV when examining the synaptic strength. Under this condition, both EPSC and IPSC exhibit inward currents. To assess the unitary synaptic transmission strength for the comparison between different groups, the postsynaptic neuron is needed to have the adequate access resistance (10–20 MΩ for PCs and PVBCs, 10–25 MΩ for ChCs) and can be well compensated. Both pre- and postsynaptic neurons are needed to be in the stable state during the recording of synaptic strength. Thus in some cases, we could confirm the presence of connectivity but were not able to measure their strength. Signals were recorded and filtered at 2 kHz, digitalized at 20 kHz (DIGIDATA 1322A, Molecular Devices) and further analyzed using the pClamp 10.3 software (Molecular Devices) for intrinsic properties and synaptic features.

#### Optical stimulation of ChR2-expressing pathways

We employed Channelrhodopsin-2(ChR2)-assisted circuit mapping^10^ to examine the local and long-range inputs to ChCs using *Nkx2.1-CreER;Ai14* mice. To investigate the local input specificity, we expressed ChR2 in a subset of PCs in PL by injecting *AAV-Ef1a-FD-ChR2-YFP* into unilateral PL and simultaneously *HSV-Ef1a-flp* into cPL or BLA, respectively. To investigate the long-range inputs, we expressed ChR2 in BLA or contralateral PL by injecting *AAV-CAG-ChR2-YFP*, respectively.

Four-eight weeks following the injection, monosynaptic responses initiated from ChR2(+) axons were recorded in postsynaptic ChCs and adjacent ChR2(−) PCs under whole-cell voltage clamp in fresh brain slice. Monosynaptic responses of these inputs were measured in the presence of 1 μM Tetrodotoxin (TTX,to block action potential generation) and 1 mM 4-Aminopyridine (4-AP, to enhance depolarization of presynaptic terminal). The laser stimulation (447 nm) was flashed (1 – 3 ms duration, usually 2 ms) on the slices through fiber LED. The laser power of the optical stimulation system at the focal plane of the slice was determined with step-wise increase of the power. We employed the power which evoked saturated responses (also see Supplementary Fig. 7).

#### In vivo Electrophysiology

##### Electrode implantation

8–10 weeks after delivering *AAV-Ef1a-FD-ChR2-YFP* or *AAV-Ef1a-FD-eYFP* into the bi-lateral PLs, 6 *Nkx2.1CreER;LSL-Flp* mice (TM induction at E17.5) were implanted with a custom made microdrive that contained electrodes and optical fibers under isoflurane anesthesia. Stereo-optrodes were implanted in the left PL (A/P: −2.00 mm, M/L: 0.20 mm, D/V: −1.18 mm). Each stereo-optrode was comprised of a 230 μm optical fiber glued to a bundle of 14 tungsten wire (13 μm diameter) stereotrodes placed 400–500 μm below the end of the optical fiber. Additionally, a 75 μm diameter tungsten wire field electrode was implanted in the ipsilateral BLA (A/P: −1.60 mm, M/L: 3.3 mm, D/V −4.4 mm). A reference screw was implanted in the skull over the frontal cortex and a ground screw in the skull over the cerebellum.

##### In vivo data acquisition

Data collection occurred 5–7 days after electrode microdrive implantation. The experiment was performed while the animal sat quiescent in a wooden box (20x3x30 cm) in the darkness. We were tracking the position of the animal at a sampling rate of 33 Hz. The laser was triggered once every second for 5 ms. In addition, there were no differences in distance traveled the ChR2 animals compared to the control animals. During the experiment there were no visible differences between the ChR2 animals and control animals upon laser stimulation. Blue light was delivered using an LED (465 nm; PlexBright LD-1 Single Channel LED Driver from Plexon). Light pulses were 5 ms long and were delivered at 1 Hz. The light power was 5 mW measured from the tip of the optical fiber patch cord. Electrophysiological data were acquired using a Digital Lynx system (Neuralynx). LFPs were referenced to a screw located in the skull over the frontal cortex / olfactory bulb, band-pass filtered (1–1,000 Hz), and acquired at 2 kHz. Single unit recordings were band-pass filtered at 600–6,000 Hz and acquired at 32 kHz; spikes were detected by thresholding and sorted off-line. Initial automated spike sorting was done based on peak, energy and the first two principal components, using Klustakwik (Ken Harris, UCL) instantiated in SpikeSort3D (Neuralynx); clusters were subsequently manually confirmed. Isolation distance and L-ratio were computed as described in^11^. The median isolation distance for the single-unit clusters was 27, and 99% of the units had an isolation distance higher than 10. The median L-ratio was 0.05, and 79% of the units had an L-ratio lower than 0.5.

##### Firing rate analysis

We used a bootstrapping analysis to determine the statistical significance of firing rate changes. First, for each cell, firing rate was binned in 5ms bins for 300 ms around each stimulus. Next, 2000 artificial “trials” were created by randomly shuffling from these bins. Distributions of firing rates were calculated for bins spanning from 0 – 15 ms (for short latency effects). Units were considered significantly modulated by the stimulus if the actual firing rates in at least three consecutive bins within either interval were below the 5th or above the 95th percentile of the shuffled distributions. This method produced very stringent (p<0.001) requirements for significance. Baseline firing rate reported in supplementary figure 10 consists of the mean firing during 200 ms prior to laser onset across all trials.

##### Phase locking and directionality PL-BLA analyses

For BLA field analyses we analyzed 3–6 Hz oscillations, as this frequency range is prominent during behavior in the BLA and has been shown to engage the PL-BLA pathway^12,13^. A given unit was said to be significantly phase locked if the distribution of the BLA LFP phases where the spikes occurred was not uniform as assessed with Rayleigh’s test for non-uniformity of circular data. Zero phase corresponds to the peak of the signal. Phase locking strength was quantified using pairwise phase consistency (PPC)^14^. Shuffled percent of phase locked cells reported in Fig. 5j were calculated by bootstrapping. Spikes were shuffled randomly 1000 and phase locking significance was calculated with rayleigh’s test and for each iteration the percent of significantly phase locked cells was calculated. On the figure we report the average percent across 1000 iterations. To calculate the power envelope and phase of BLA theta, a bandpass filter for 3–6 Hz was used using a zero-phase-delay FIR filter with Hamming window (filter0, provided by K. Harris and G. Buzsaki, New York University, USA), the phase component was calculated by a Hilbert transform, and a corresponding phase was assigned to each spike. To analyze the directionality of PL phase-locking to BLA theta (3–6 Hz), single units with at least 50 spikes were included because the MRL statistic can be highly variable for small spike numbers. The LFP times were lagged relative to the spike timing from −100 ms to 100 ms, stepping by 5 ms, and the MRL value was determined for each single unit at each lag. The MRL at each lag was normalized by dividing by the mean MRL across all lags. For the BLA evoked potential analysis, the BLA LFP was average across all trials (5 ms light presentations), for each animal and the mean evoked potential for 5–10ms post light presentation was compared across ChR2 and eYFP injected mice with two sample t-test.

#### Analysis and Statistics

For comparison of two groups of data, Mann-Whitney test and Student’s paired t-test and two-sample t-test were used as indicated in the experiment. To compare the observed distributions, Pearson Chi-Square tests (Fisher exact tests if some number of samples below 5) were used. In cases where statistical differences were assessed between brain regions with rabies traced input sources, one-way ANOVAs were performed followed by Tukey-Kramer tests for mean comparisons. Data are presented as mean ± s.e.m. if not specifically indicated, and a p value <0.05 was considered significant. The significance was marked as *, p < 0.05; **, p <0.01 and ***, p < 0.001.

## Supplementary Figures and Table

**Supplementary Figure 1.**
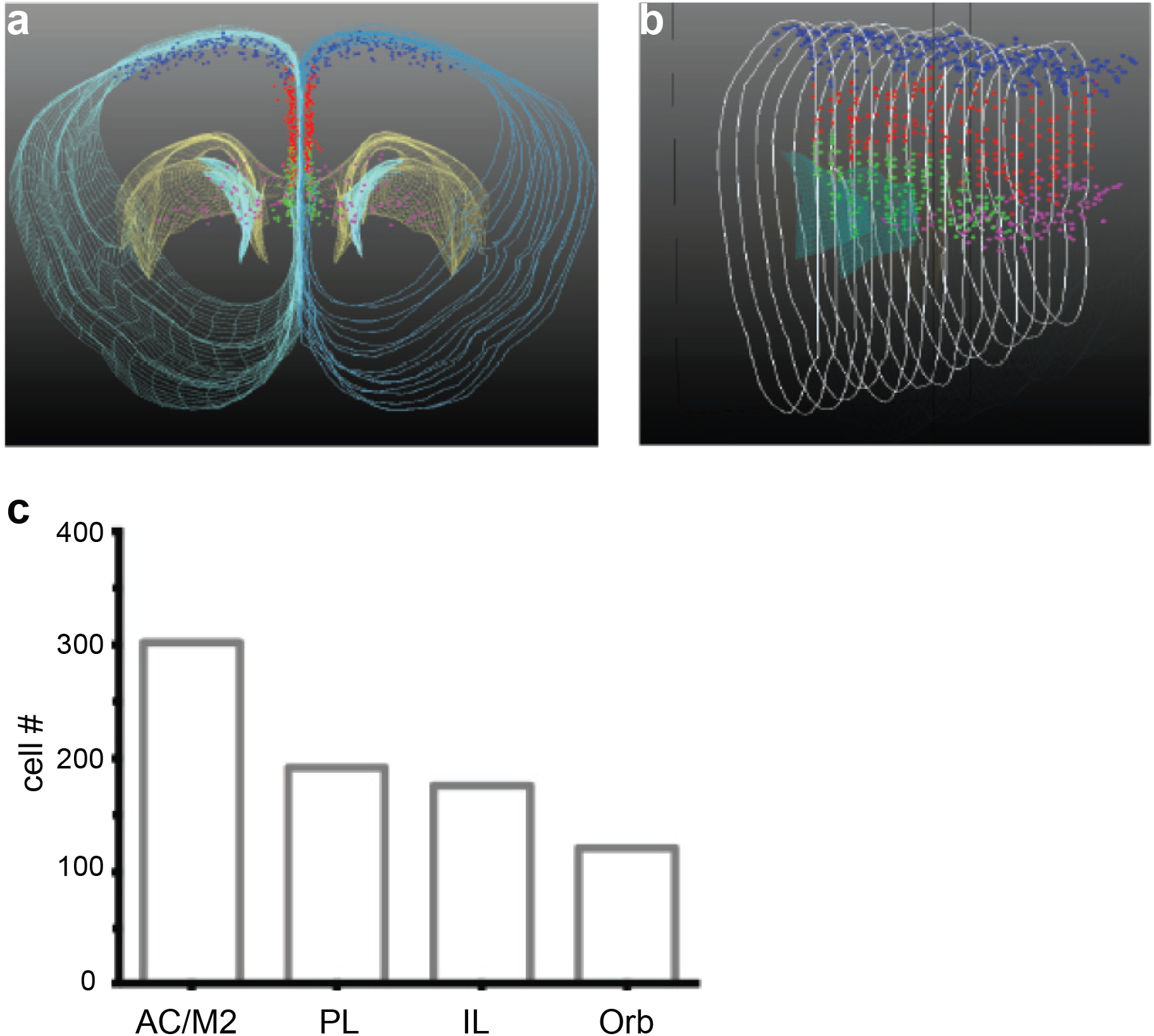
Spatial distribution of L2 ChCs in frontal cortex. (**a-b**) ChCs were labeled by tomaxifen induction at E17.5 in a *Nkx2.1-CreER;Ai14* mouse. Images were taken from 100 um coronal sections, and stereologically reconstructed across 1 mm thickness of cortical tissue. Green dots are in infralimbic cortex (IL), red dots in prelimbic cortex (PL), blue dots in anterior cingulate cortex and secondary motor cortex (AC/M2), and purple dots in orbitofrontal cortex (Orb). (**c**) Quantification of ChCs labeled in several subregions of frontal cortex in a single animal.

**Supplementary Figure 2.**
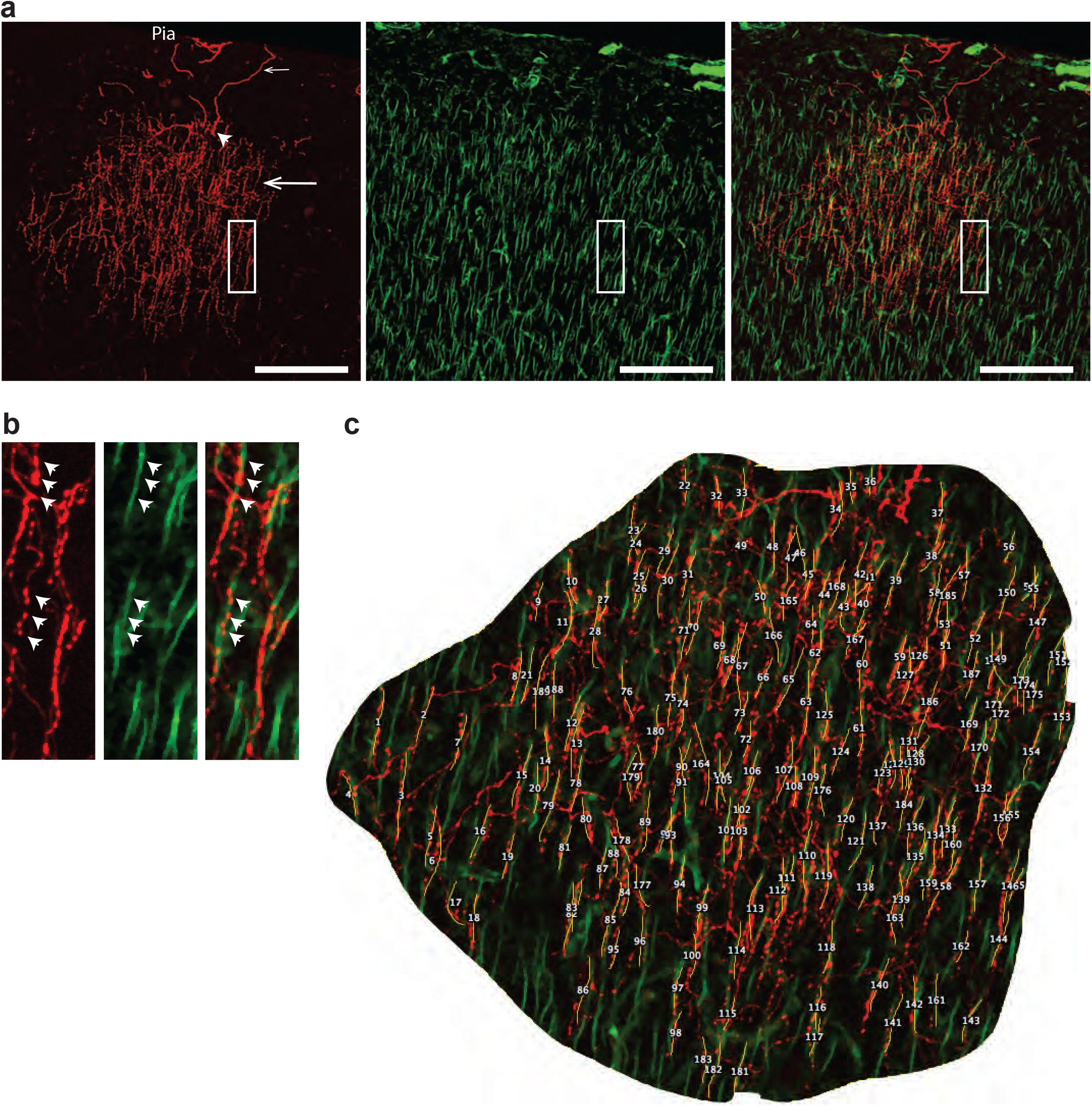
Counting of axon cartridge number of individual L2 ChCs in PL. (**a**) Left: a RFP(+) L2 ChC from spare labeling with lose dose TM induction in Nkx2.1-CreER;Ai14 (arrowhead: soma, small arrow: dendrite, larger arrow: axon arbor). Middle: staining of PC AIS with phosphor-IkappB. Right: overlay. Arrows indicate soma and dendrite position. (**b**) The cartridges revealed as strings ChC axon buttons (arrowheads) along the AIS. Images are magnified from boxes in (**a**). (**c**) Example of counting cartridge number of a ChC.

**Supplementary Figure 3.**
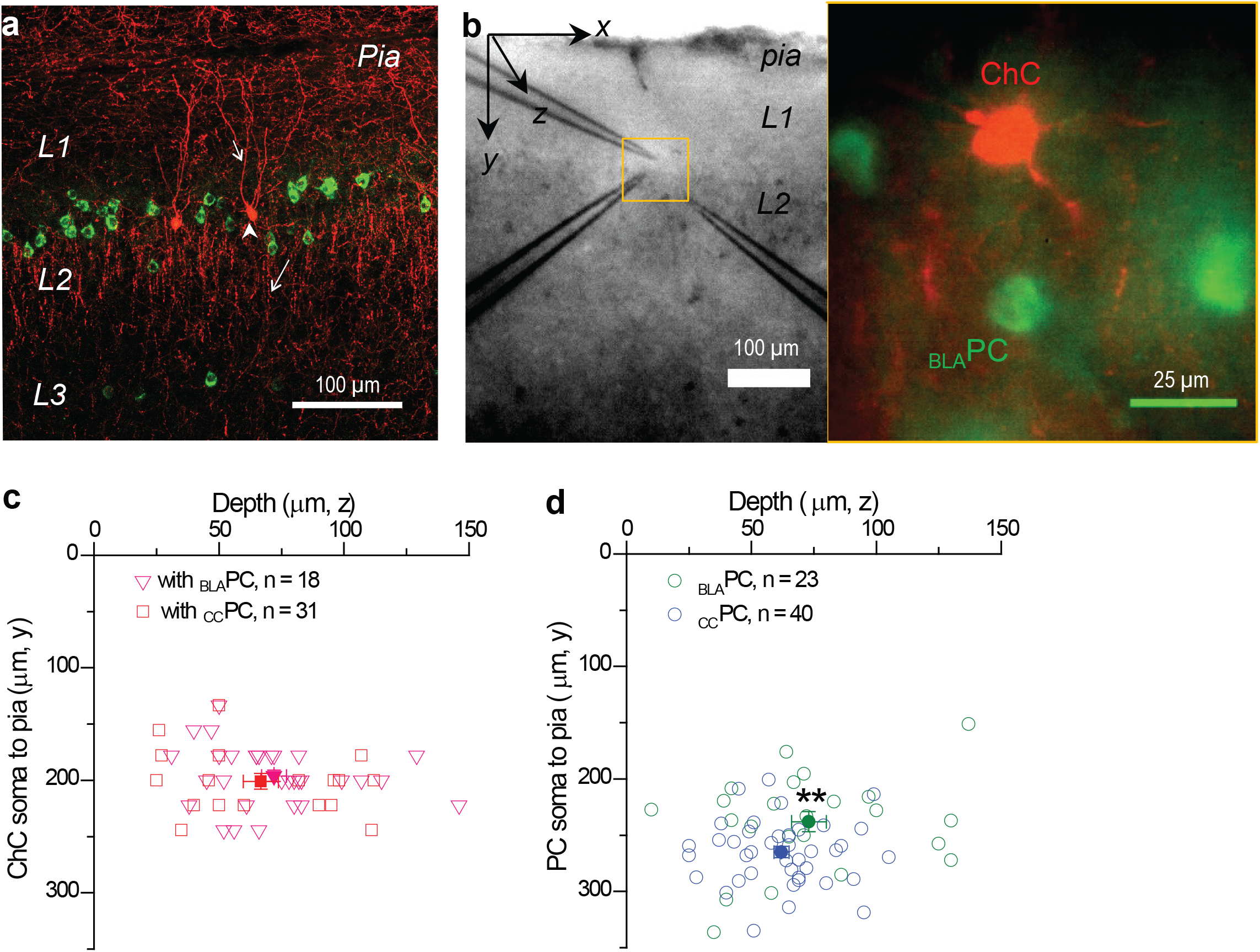
Spatial relationship between ChCs and PCs in PL and in paired patch recording experiments. (**a**) RFP(+) L2 ChCs in PL were positioned alongside to CTB-488 labeled _BLA_PCs. Dendrites (short arrow), soma (arrowhead) and axons (long arrow) of ChCs are indicated. (**b**) An example of whole-cell patch recordings of a L2 ChC and an adjacent CTB labeled _BLA_PC in brain slice. Right panel of pseudocolored cells taken from boxed area of the left under fluorescent imaging. (**c**) Position of all recorded ChCs in brain slice with dimension axis as indicated in b. In the distance from their somata to pia (201±7 μm vs.196±5 μm, p=0.33, Mann-Whitney test). In the depth in the slice (z axis) (67±7 μm vs 72±5 μm, p=0.46, Mann-Whitney test). (**d**) Position of all recorded _BLA_PCs and _CC_PCs in the brain slice. In the depth (z axis) (62±3 μm vs 73±7 μm, p=0.33, Mann-Whitney test). Mann-Whitney test found a significant difference (p<0.01) in the distance from their somata to pia (238±9 μm vs 265±5 μm).

**Supplementary Figure 4.**
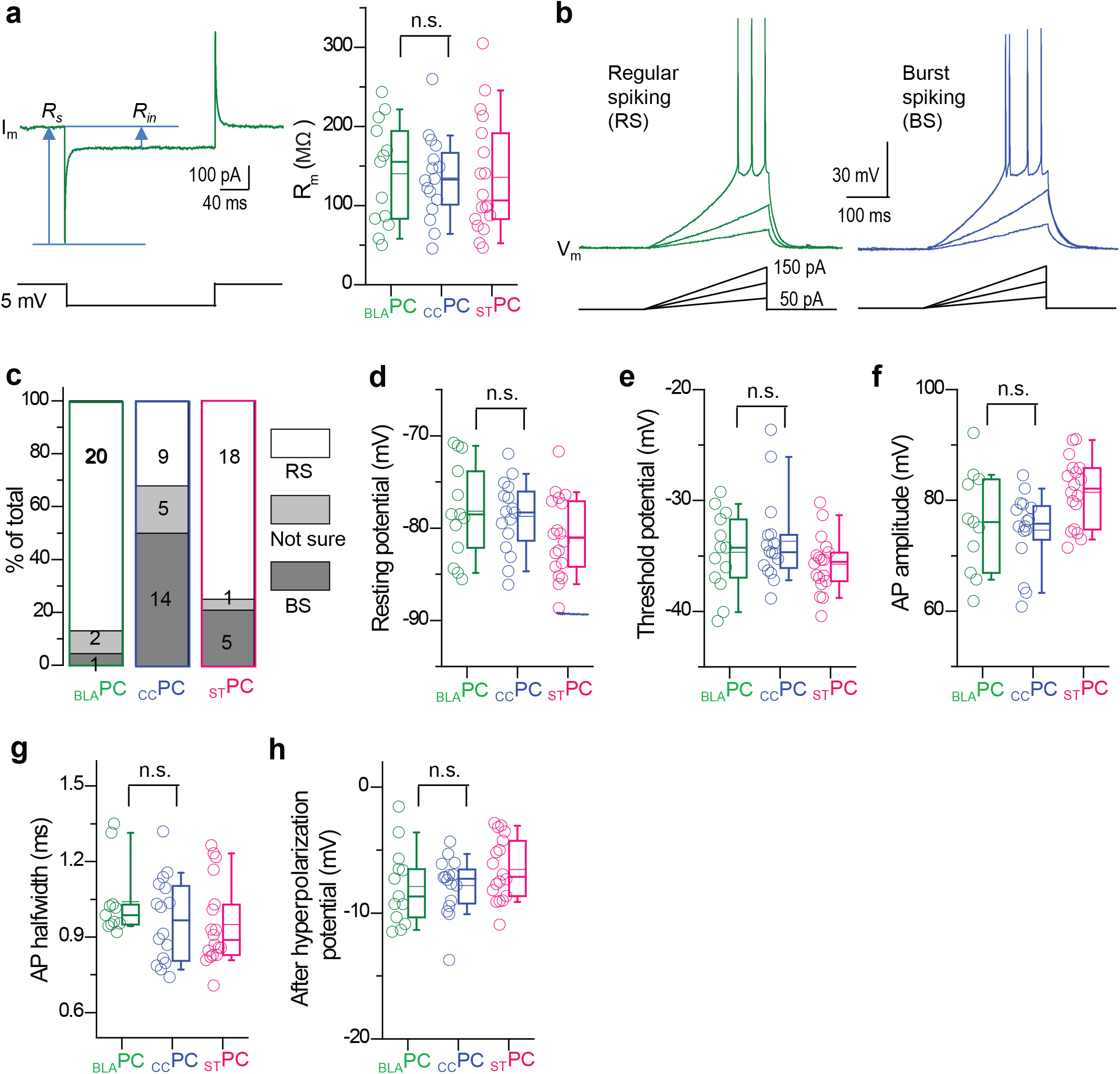
Intrinsic properties of 3 subsets of PCs in the upper layer of PL. (**a**) An example of membrane resistance (R_m_) measurement by a 5 mV hyperpolarization under voltage-clamp; summary in right panel. No significant difference (p=0.81, Mann-Whitney test) between _BLA_PCs (140±66 MO, n=13) and _CC_PC (135±53 MΩ, n=16). **(b**) Examples of firing properties showing regular spiking (RS) and burst spiking (BS) in two PCs under current-clamp. Step-wise ramp depolarization currents were injected into the soma through the glass pipettes. BS was recognized as an interval (<10 ms) between initial two spikes. (**c**) The percentage of RS and BS in each subset of PCs. (**d-h**) Summaries of resting membrane potential, AP threshold, AP amplitude, AP half-width and after-hyperpolarization potential of 3 subsets of PCs. _BLA_PCs (n=13), _CC_PCs (n=16) and _ST_PCs (n=18). Comparison of _BLA_PCs vs _CC_PC by Mann-Whitney test showed no significant difference in resting membrane potential (−78.2±5.2 vs −78.7±3.9 mV, p=0.88), AP threshold (−34.7±3.6 vs −33.7±3.9 mV, p=0.74), AP amplitude (76.1±9.2 vs 74.6±6.7 mV, p=0.67), AP half-width (1.04±0.15 vs 0.97±0.17 ms, p=0.42), and after-hyperpolarization potential (7.9±3.0 vs 7.8±2.3 mV, p=0.71). Values are presented as mean±s.d.. Plots display median, mean, quartiles and range.

**Supplementary Figure 5.**
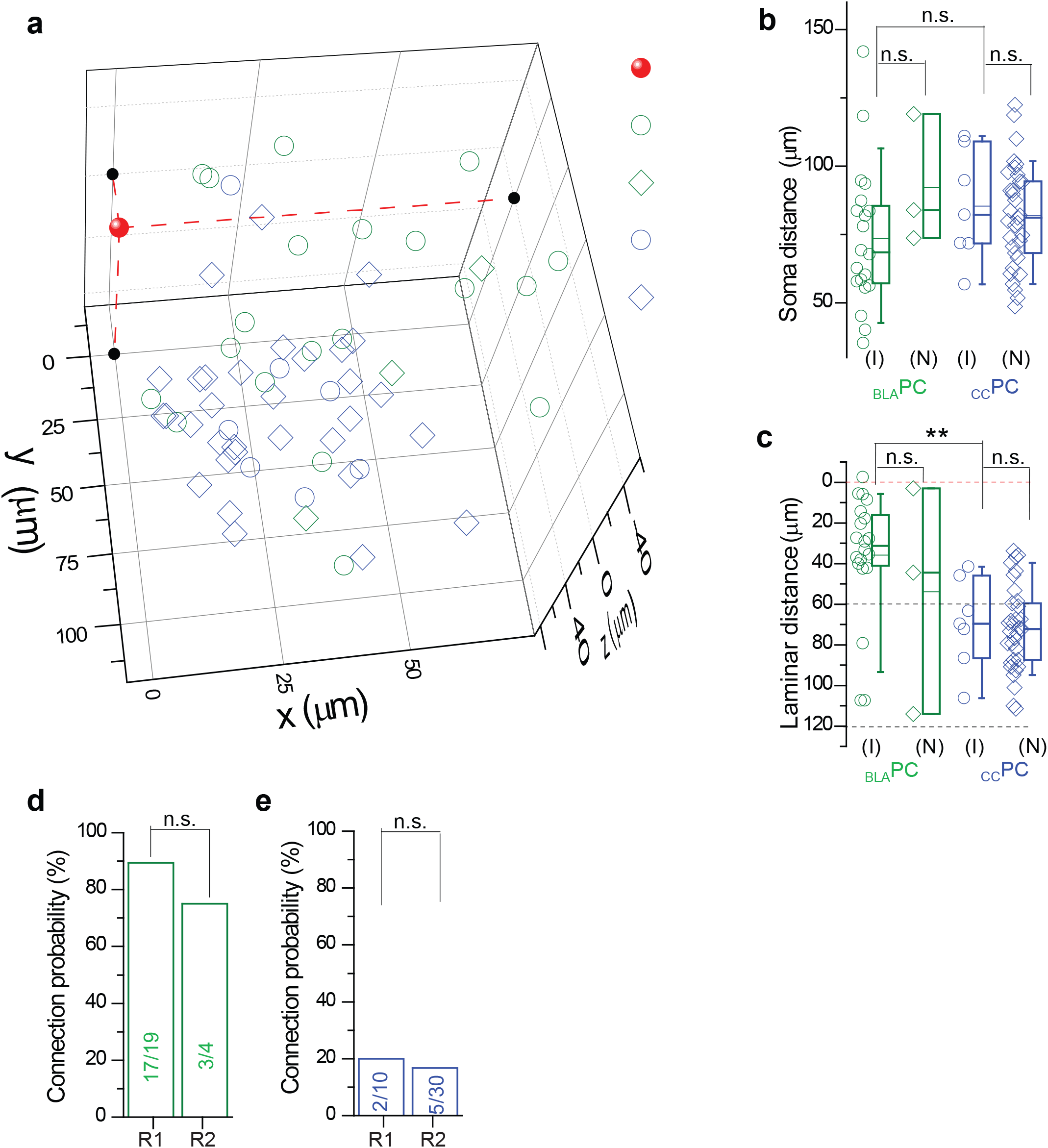
Quantification of spatial relationship between L2 ChCs and PCs in paired recording experiments. (**a**) 3D plots of all recorded PCs aligned to the recorded ChCs as origin (the red dot) of the 3D axis. Innervated (I) _BLA_PCs: green circle, n=20; innervated _CC_PCs: blue circle, n=7; non-innervated (N) _BLA_PCs: green diamond, n=3; non-innervated _CC_PCs: blue diamond, n=33. (**b**) Summarizes of soma distance to ChC for the four categories of PCs in (**a**). No significant difference was detected in distance to ChCs between I-_BLA_PCs vs. N-_BLA_PCs (73±26 μm vs 92±24 μm, p=0.19), or between I-_CC_PCs vs. N-_CC_PCs (85±20 μm vs 82±19 μm, p=0.70), or between I-_BLA_PC vs. I-_CC_PC (p=0.19). (**c**) Summarizes of laminar distance to ChC for the four categories of PCs. There was no significant difference detected between I-_BLA_PCs vs. N-_BLA_PCs (35±30 μm vs 54±26 μm, p=0.49) or between I-_CC_PCs vs. N-_CC_PCs (69±22 μm vs 72±21 μm, p=0.72). In the laminar distance to ChCs, I-_BLA_PCs are significantly closer than I-_CC_PCs (p<0.01). Values are presented as mean±s.d. and Mann-Whitney test employed. (**d-e**) Summary showing that the connection probability (numbers in bars indicate connected / tested pairs) from ChCs to _BLA_PCs (**d**) and _CC_PCs (**e**) were the same whether PCs were located 0-60 μm (R1) or 60-120 μm (R2) from ChCs. Although the majority of _BLA_PCs were located in the upper L2 close to ChCs (R1: n=17 out of 19 pairs, 89.5 %), the more sparse _BLA_PCs in deeper L2/3 were innervated by L2 ChCs with similar probability (R2: n=3 out of 4 pairs, 75.0 %; Fisher exact test, p=0.45). L2 ChCs had similarly low innervation probability (Fisher exact test, p = 0.99) for _CC_PCs located in the upper (R1: n=2 out of 10 pairs, 20.0 %) or deeper (R2: n=5 out of 30 pairs, 16.7 %) L2/3. Plots display median, mean, quartiles and the range.

**Supplementary Figure 6.**
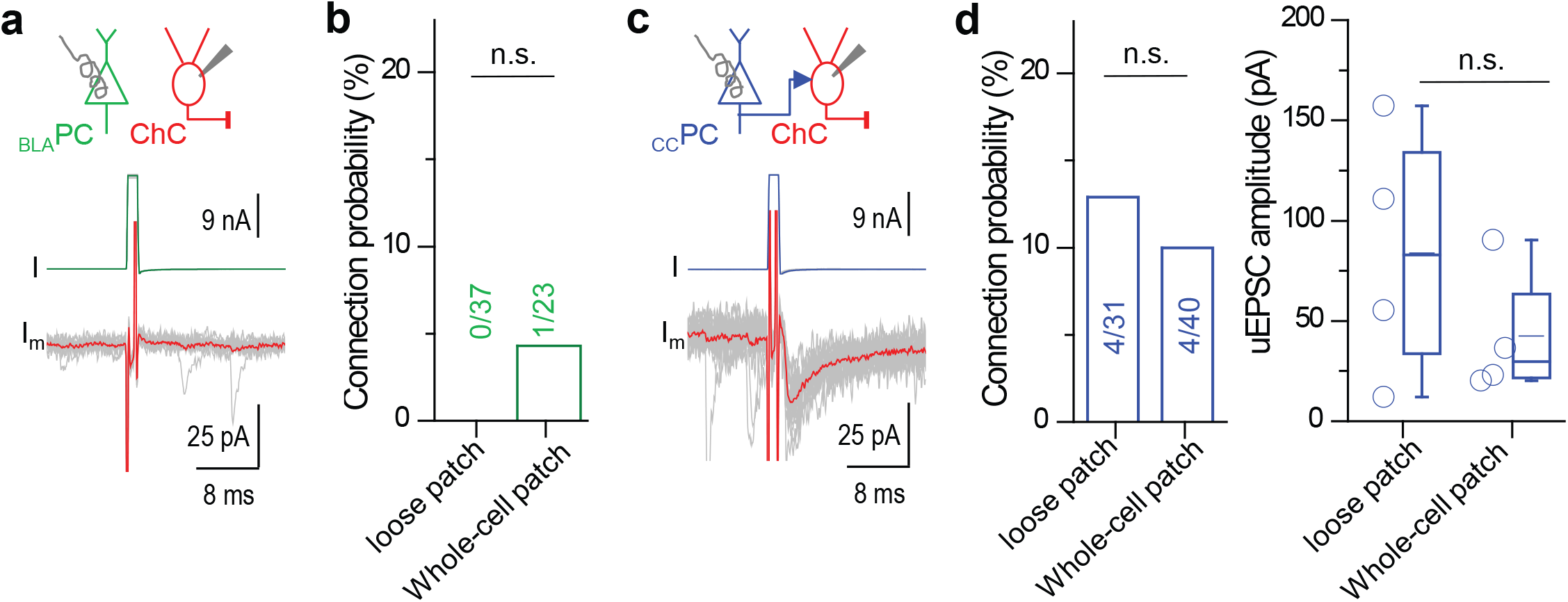
Detection and quantification of PC to ChC connection by loose patch stimulation of PCs. Loose patch was achieved by the tight seal (> 100 MΩ resistance) to the targeted PCs through the same pipette for the whole-cell patch recordings. Stimulation ranged from 0.1-1 V in 200 μs at 0.1Hz. Stimulation intensity was determined by the persistence of spikes in PCs after the stimulation artifact. (**a**) Example of lack of synaptic responses in ChCs (red) following loose patch stimulation of a _BLA_PC (green). Upper: schematic of loose patch stimulation in _BLA_PC and wholecell patch recording in ChC. Lower: representative traces from paired recordings in ChCs; thick trace was averaged from 20 trials. (**b**) Summary of _BLA_PC to ChC connection probability (numbers in bar graph indicate connected / tested pairs). Fisher exact test revealed no significant difference (p=0.38) in the connection probability by loose patch stimulation of _BLA_PCs (n=0 out of 37 pairs, 0.0 %) compared to that by whole cell patch stimulation of _BLA_PCs (n=1 out of 23 pairs, 4.3 %). Thus, we pooled two sets of data for the analysis in Fig. 2c. (**c**) Same arrangement as in a showing example of synaptic responses in ChCs (red) following loose patch stimulation in _CC_PC (blue). (**d**) Same arrangement as in b showing summaries of _CC_PC to ChC connection probability (numbers in bar graph indicate connected / tested pairs). Fisher exact test revealed no significant difference (p=0.72) in the connection probability by loose patch stimulation of _CC_PCs (n=4 out of 31 pairs, 12.6 %) compared to that by whole cell patch stimulation (n = 4 out of 40 pairs, 10.0 %). Thus, we pooled two sets of data for the analysis in Fig. 2c. Mann-Whitney test revealed no significant difference (p=0.49) in the synaptic strength evoked by loose patch stimulation (83.9±31.7 pA, n=4) and by whole cell patch stimulation (42.5±16.3 pA, n= 4). Thus, we pooled two sets of experiments together for the analysis in Fig. 2c. Plots display ttmedian, mean, quartiles and the range.

**Supplementary Figure 7.**
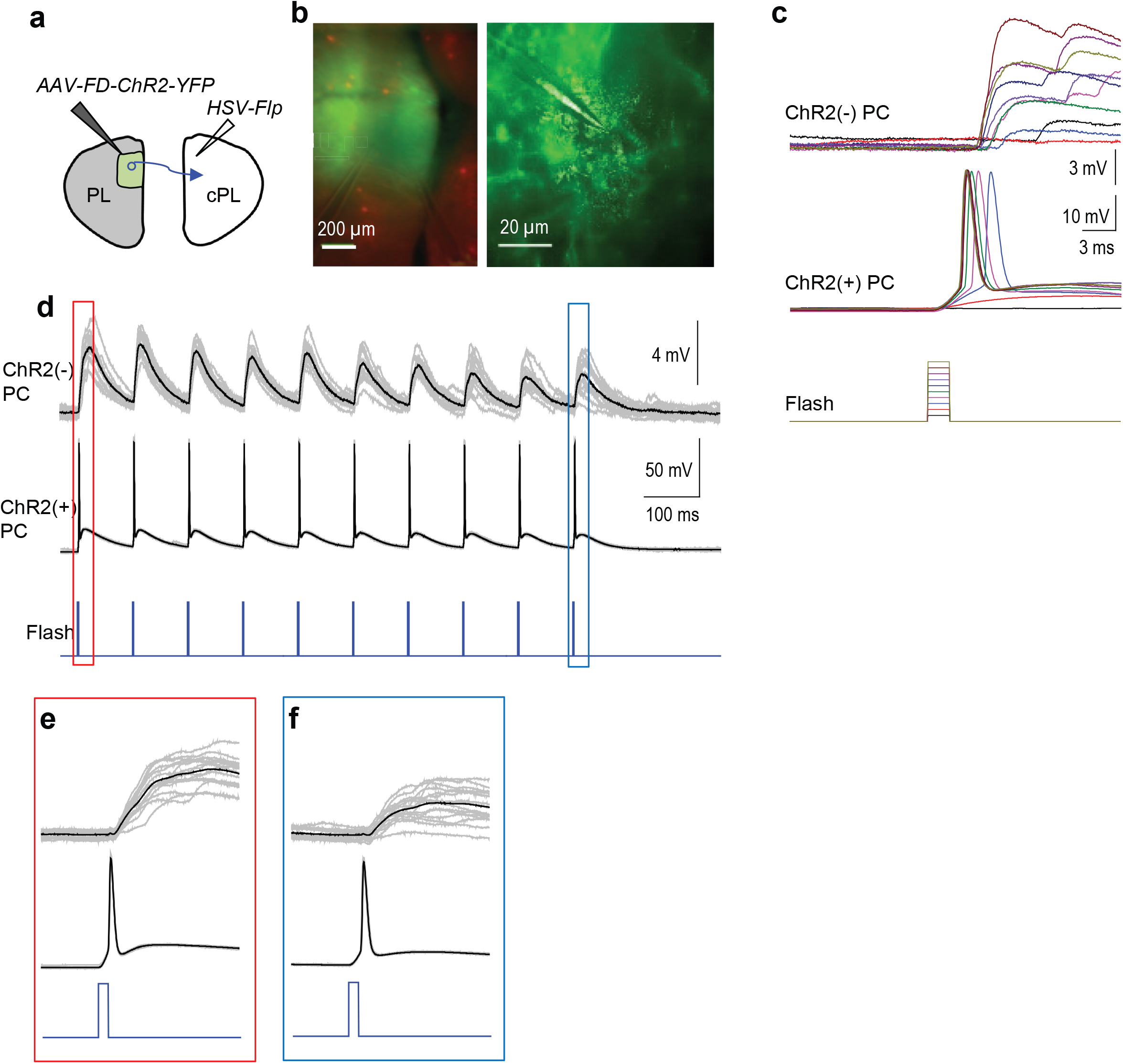
Reliable light-induced spiking in ChR2(+) _CC_PCs and postsynaptic responses in adjacent ChR2(−) PCs. (**a**) Schematic _CC_PCs infection by dual viral injections first with *HSV-Flp* in cPL and then with *AAV-FD-ChR2-YFP* into PL. (**b**) Example paired recording of YFP(+) PC and adjacent YFP(−) PC in cortical slice under fluorescent microscope. Red: RFP(+) L2 ChCs; Green: ChR2(+) _CC_PCs in PL. (**c**) Example spikes in a ChR2(+) evoked by step-wise increase of blue laser intensity (2 ms duration) and postsynaptic potentials in a ChR2(−) PC. (**d**) Example of high fidelity of spiking in a ChR2(+) PC and synaptic potential in a ChR2(−) PC (black trace averaged from 15 trials of gray traces) evoked by laser flash of 2 ms duration at 10 Hz with a saturation intensity based on step-wise measurement in **c**. (**e**) and (**f**) were magnified from the corresponding events indicated in **d**.

**Supplementary Figure 8.**
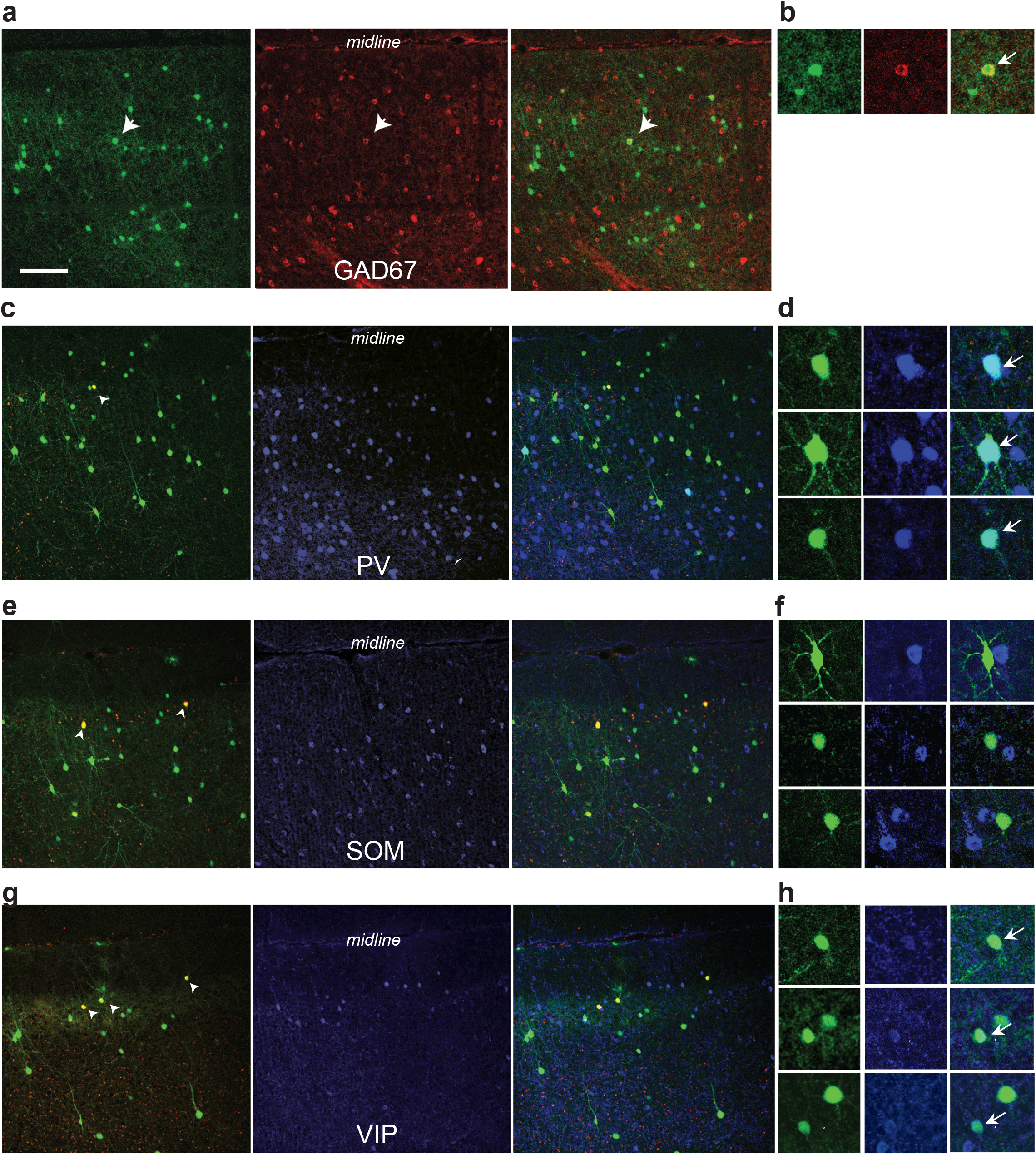
Identification of GABAergic inputs to L2 ChCs revealed by rabies tracing. (**a**) Left: local EnvA-dG-GFP infected cells. Middle: GAD67 immunohistochemistry. Right: overlay. (**b**) Incidence of colocalized inputs with GAD67. Indicated by the arrow as in **a**. (**c**) Left: local EnvA-dG-GFP infection RFP+ starter ChCs are designated by white arrow head. Middle: PV immunohistochemistry. Right: overlay. (**d**) Incidence of colocalized inputs with PV as indicated by arrows. (**e, f**) Same as in (**c, d**) instead stained for somastain (SOM), note absence of colocalized input. (**g, h**) Same as in (**c, d**) instead stained for vasoactive intestinal peptide (VIP). Scale bar: 100 μm.

**Supplementary Figure 9.**
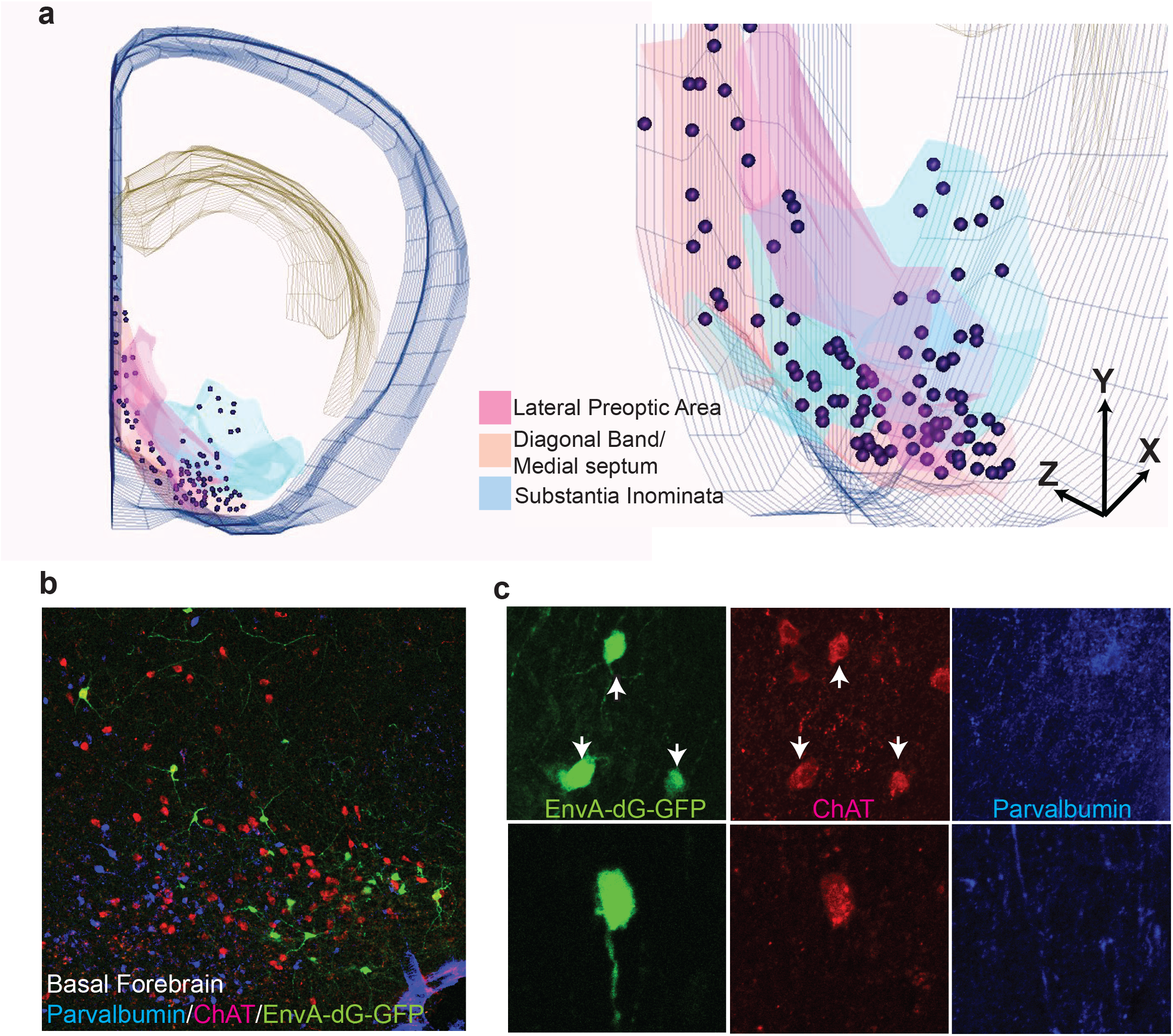
Identification of diagonal band (DB) input to L2 ChCs revealed by rabies tracing. (**a**) Left: 3D reconstruction for total basal forebrain inputs from a single animal color-coded by individual nuclei. Right: 45 degree rotation zoomed image of that basal forebrain. (**b**) Basal forebrain immunostained for PV (blue) and ChAT (red), overlaid with *EnvA-dG-GFP* rabies labeled inputs to L2 ChCs in PL. (**c**) Incidence of inputs colocalized with ChAT (arrowheads) but negative for PV.

**Supplementary Figure 10.**
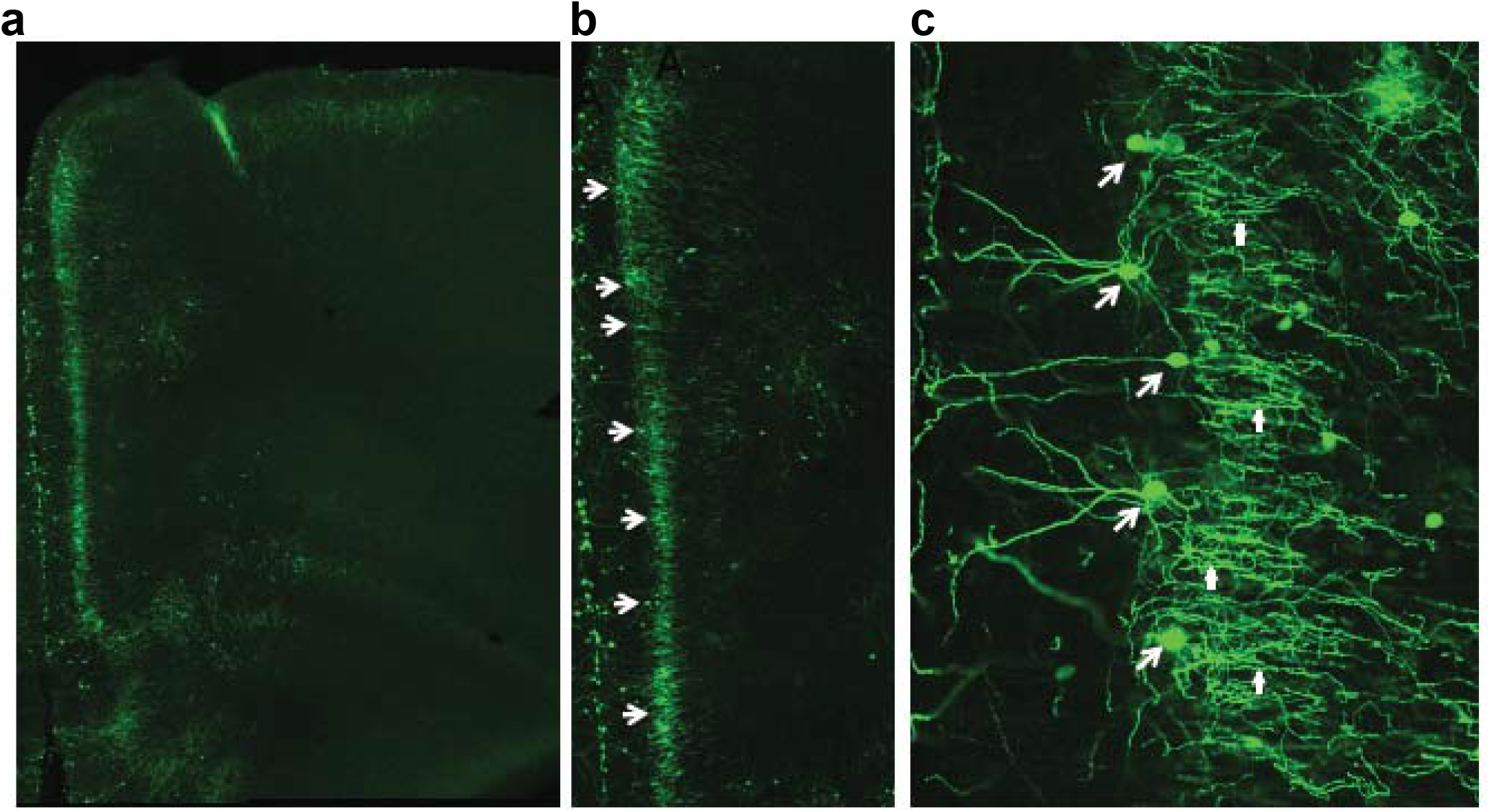
The specificity of mPFC ChCs labeling by the injection of *AAV-FD-ChR2-YFP* in *Nkx2.1CreER;LSL-Flp* mice induced at E17.5. (**a**) Injection site following injection with *AAV-FD-ChR2-YFP* into mPFC, amplified and co-stained with antibodies against YFP tag. (**b**) Zoomed in injection site reveals several morphologically identified ChCs (indicated by arrows). (**c**) Morphologically distinct and homogenous populations ChCs can be identified via dendritic trees in layer 1, somata on the layer 1/2 border (large white arrows) and extensive axonal cartridge plexus in layer 2/3 (small white arrows). Of total viral label cells, 88.6± 5.6 % cells were ChCs and in layer 2/3 specifically 94.8±3.6% of cells were ChCs (n=5 animals). No incidences of virally labeled pyramidal cells were seen.

**Supplementary Figure 11.**
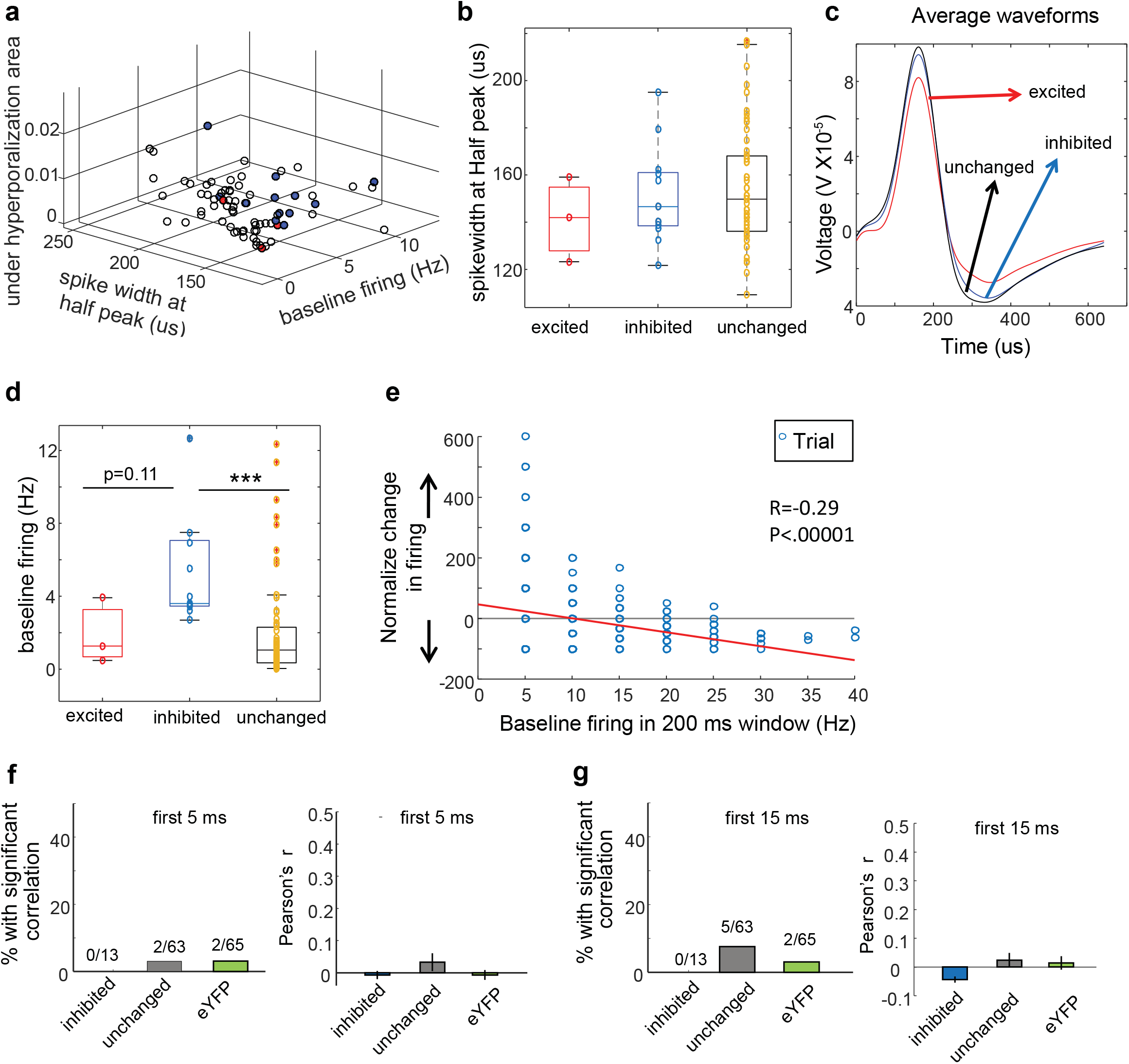
(**a**) Plot showing hyperporalization, spike width and baseline firing for all recorded units identified by their effects in firing by ChC activation. The subpopulation of units indicated by colors as showing in **b-d**. Differences in spike width (**b**), average waveform (**c**) and baseline firing (**d**) for the different subpopulations of single units recorded. (excited n=3; inhibited n=13; unchanged n=63; Mann Whitney test inhibited vs unchanged: ***p<.001; z-score: 4.2626). (**e**) Example trial by trial correlation between baseline firing and ChC activation induced changes in firing in one single unit. (**f-g**) Across the inhibited, excited and unchanged populations at different time windows post laser: 0–5 ms (**f**), and 0–15 ms (**g**) on the left, the percent of cells with a significant correlation between the baseline firing rate and the firing rate effect of ChC activation (no proportion is statistically different from eYFP group) and on the right, the correlation coefficients for the inhibited and unchange subpopulations are not different than those for control group (inhibited n=13; no change n=63; eYFP n=65; Mann Whitney test, inhibited vs eYFP p=0.32; unchange vs eYFP p=0.85, in 0-15 ms window).

**Supplementary Figure 12.**
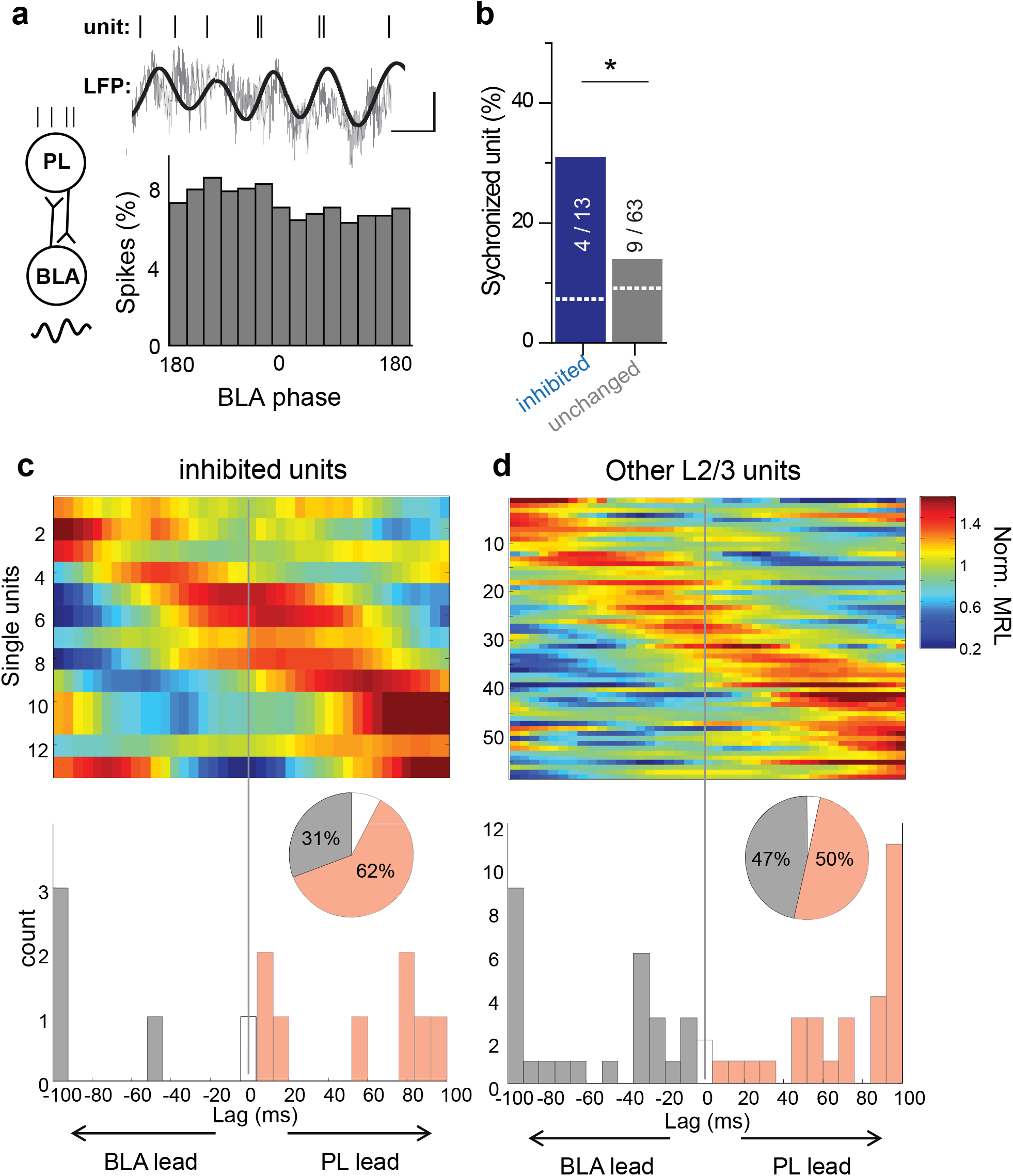
(**a**) Phase-locking between PL single units and BLA LFP, depicted in the left schematic. Top: An example of phases locking: spikes from a PL unit appeared to occur at or near the peaks of the raw (gray) and filtered (black) BLA LFP (top); bottom: PSTH and phase distributions of the representative PL single units. Bin size in all PSTHs is 2 ms. (**b**) Summary of the percentage of phase-locked units separated into light-inhibited and unchanged groups (numbers in bars indicate phase-locked / total units in the group). Significance of phase-locking was determined by Rayleigh’s test. The dashed lines indicated the amount of phase locking with shuffled data. The probability of obtaining as many phase-locked units as were seen in the actual dataset were p<0.01 for the inhibited units, and p=0.17 for the non-inhibited units (p values were obtained with bootstrapping methods). Light-inhibited units were preferentially synchronized to BLA LFP. (**c-d**) Assessing PL-BLA directionality with a lag analysis. Top, phase-locking strength to BLA LFP 3–6 Hz measured as normalized mean resultant length (MRL) as a function of lag for inhibited units (**c**) and other L2/3 units (**d**) aligned by peak lag. Bottom, distributions of lags at which peak phase-locking occurred, for inhibited units (**c**) and for other L2/3 units (**d**). Inset pie charts show in gray units with negative peak lags, in red units with positive peak lags, and in white units with a lag of zero. Units had to have at least 50 spikes to be included in these analyses.

## Lu&Tucciarone-NN-Supplementary Table 1

**Table 1.**
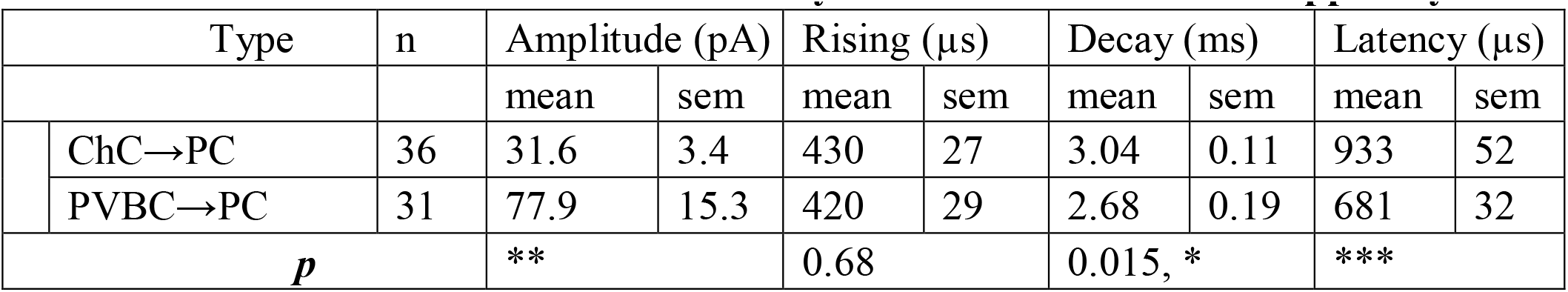
Kinetics of uIPSC in PCs innervated by ChCs and PVBCs in PL upper layers.

